# Phylogenetic Characterization of the Energy-taxis Receptor Aer in *Pseudomonas* and Phenotypic Characterization in *P. pseudoalcaligenes* KF707

**DOI:** 10.1101/081455

**Authors:** Sean C. Booth, Raymond J. Turner

**Author notes:** Address correspondence to Raymond J. Turner, and Sean C. Booth. Department of Zoology, Department of Biochemistry, University of Oxford, UK.

## Abstract

Chemotaxis allows bacteria to sense gradients in their environment and respond by directing their swimming. Aer is a receptor that, instead of responding to a specific chemoattractant, allows bacteria to sense cellular energy levels and move towards favourable environments. In *Pseudomonas*, the number of apparent Aer homologs differs between the only two species it had been characterized in, *P. aeruginosa* and *P. putida*. Here we combined bioinformatic approaches with deletional mutagenesis in *P. pseudoalcaligenes* KF707 to further characterize Aer. It was determined that the number of Aer homologs varies between 0-4 throughout the *Pseudomonas* genus, and they were phylogenetically classified into 5 subgroups. We also used sequence analysis to show that these homologous receptors differ in their HAMP signal transduction domains. Genetic analysis also indicated that some Aer homologs have likely been subject to horizontal transfer. *P. pseudoalcaligenes* KF707 was unique among species for having three Aer homologs as well as the receptors CttP and McpB. Phenotypic characterization in this species showed the most prevalent homolog of Aer was key, but not essential for energy-taxis. This study demonstrates that energy-taxis in *Pseudomonas* varies between species and provides a new naming convention and associated phylogenetic details for Aer chemoreceptors.

## Introduction

Chemotaxis is the ability to sense and swim along chemical gradients, and is a widespread, important behaviour in bacteria [1]. Canonically, it functions by extracellular compounds interacting with membrane-bound chemoreceptors, generally called methyl-accepting chemotaxis proteins (MCPs), causing a phosphorylation signal cascade through CheA and CheY to alter the direction of flagellar rotation [1], which in turn allows the cell to direct swimming through concentration gradients. The first chemoreceptors were characterized in *E. coli*, but now many more have been described, particularly in *Pseudomonas* [2]. Many receptors for specific ligands are being identified through a high-throughput approach [3], but some specialized receptors do not recognize extracellular signals.

Energy-taxis is a behavior in bacteria that enables swimming towards optimal environments for producing energy by sensing intracellular signals [4]. This is closely related to aerotaxis (swimming towards oxygen) as in *E. coli*, a receptor was discovered which enabled taxis towards air bubbles [5]. This inner membrane receptor was named ‘Aer’ though it does not bind oxygen, but instead enables taxis towards metabolizable carbon sources [6] by detecting the cellular energy state via a cytosolic PAS (Per-Arnt-Sim) domain with a flavin adenine dinucleotide (FAD) co-factor [7]. A similar receptor was also discovered in *P. aeruginosa*, along with a second distinct protein, named Aer-2 as both contributed to aerotaxis [8]. Aer-2 is a cytosolic protein that binds oxygen through a heme group [9]. In *P. putida* three *aer*-like genes were found, but only one was important for energy-taxis [10]. This was also named ‘Aer2’ so we will refer to the cytosolic receptor Aer-2 as McpB hereafter. It has been shown to bind oxygen, but does not affect flagellar activity as it is key to the assembly of, and signals through the Che2 chemosensory system (whose function is unknown) [11–13].

Aer has been characterized by gene inactivation in *P. putida* PRS2000 [14], *P. aeruginosa* PA01 [8], *P. putida* KT2440 [10] and *P. putida* F1 [15]. The *P. putida* species all have 3 *aer*-like genes but not *mcpB*, whereas *P. aeruginosa* only has one, indicating there is variation within the genus for both loci. The species our group studies, *P. pseudoalcaligenes* KF707, has 3 *aer*-like genes despite being more closely related to *P. aeruginosa* than *P. putida* [16, 17] and three chemosensory clusters, corresponding to the *P. aeruginosa* F6, F7 and pil-chp systems [18], making it possible to investigate the function of multiple *aer*-like genes and *mcpB* in the same organism. In *P. aeruginosa*, McpB was originally implicated with Aer as an aerotaxis receptor [8], but its function remains uncertain as despite its ability to directly bind oxygen, its signal is not transduced to the flagellum [9]. The *mcpB* gene is part of the *che2* gene cluster, which is preceded by another chemoreceptor, *mcpA* [13]. This gene was demonstrated to mediate positive chemotaxis towards tetrachloroethylene and renamed *cttP* [19]. As CttP has no obvious ligand-binding region and in *E. coli*, its MCPs can mediate attractant and repellent responses to phenol that do not involve their ligand binding regions [20]; the observed taxis to TCE may also have been fortuitous, indicating that CttP has some other function. Additionally, CttP co-localizes with the che (F7) chemosensory system while McpB is necessary for the formation of the che2 (F6) chemosensory complex [13] which led us to question whether CttP and McpB may somehow influence energy-taxis.

In this study we present a combination of a bioinformatic characterization of Aer throughout the *Pseudomonas* genus with a genetic knockout characterization of three Aer homologs, McpB and CttP in *P. pseudoalcaligenes* KF707. A phylogeny of Aer was built using sequences obtained from 65 *Pseudomonas* species providing insight into the distribution throughout the genus and enabling the definition of five ‘Aer’ groups. Only *P. pseudoalcaligenes* KF707 had the unique feature of possessing three Aer homologs, McpB and CttP making it the ideal candidate to investigate if these receptors had related functions. Using single and combinatory deletion mutants, we found that the most common *Pseudomonas* Aer homolog had the most influence on energy-taxis. Together these results provide a definition of the Aer energy-taxis receptor as a family with varied distribution in *Pseudomonas* and implicate the most common Aer homolog as the primary energy-taxis receptor in *P. pseudoalcaligenes* KF707.

## Results

To investigate how the number of Aer-like receptors, differences in their amino acid sequences and how they are related phylogenetically, Aer sequences from 65 *Pseudomonas* species were analyzed. Species with completely sequenced genomes were selected, but for those with many strains only a few representatives that have been highly studied were included (e.g. *P. aeruginosa*). Species with incomplete (draft) sequences were also included in an attempt to ensure representation from all major *Pseudomonas* clades [16, 17]. 144 protein sequences were obtained from the NCBI database using the *P. aeruginosa* PA01 Aer sequence (NP_250252.1) as a BLAST [21] query sequence. All hits with >95% sequence coverage were included. These sequences were aligned using COBALT [22] as this alignment algorithm ensures that conserved domains are aligned despite a lack of similarity elsewhere in the sequence.

From this alignment a maximum likelihood phylogeny was constructed using PHYML (Supplementary Figure 1) [23, 24]. Two additional phylogenies were generated, one including Aer from *E. coli* K12, and one including seventeen Aer-like sequences from the eight closest related species to *Pseudomonas* [25]. The topology of these two unrooted phylogenies (Supplementary Figures 2 and 3) were identical to the *Pseudomonas*-only tree, which was rooted based on the similar placement of the outgroups in both trees (Supplementary Figure 4). The rooted phylogeny allowed clear delineation of Aer sequences into subgroups.

### Phylogenetic Grouping of Aer

The generated phylogeny implied that there are several sub-families of Aer homologs (Figure 1B). These groups were sub-aligned by themselves, then analyzed to determine if the group assignments were accurate. Sequence Harmony/Multi Relief [26] was used to determine, for each position, whether that AA is conserved within each group, and whether it is divergent between the two groups (Supplementary Figure 5). Each pair of groups was compared in this fashion and most intergroup comparisons indicated that 25-35% of AAs were above the distinction cut-off, indicating the group distinctions were correct. Conversely, two proposed subgroupings of Aer.g1 were demonstrated to differ from the rest of Aer.g1 by much less than any other between-group comparisons as only 10% of AAs were above the cut-off. These two groups were thus included as part of Aer.g1 (named Aer.g1A and Aer.g1B), reducing the total number of groups to five. Throughout this process, Aer.g5 was consistently excluded as it only contains 2 sequences, which are highly divergent from all others.

**Figure 1:**
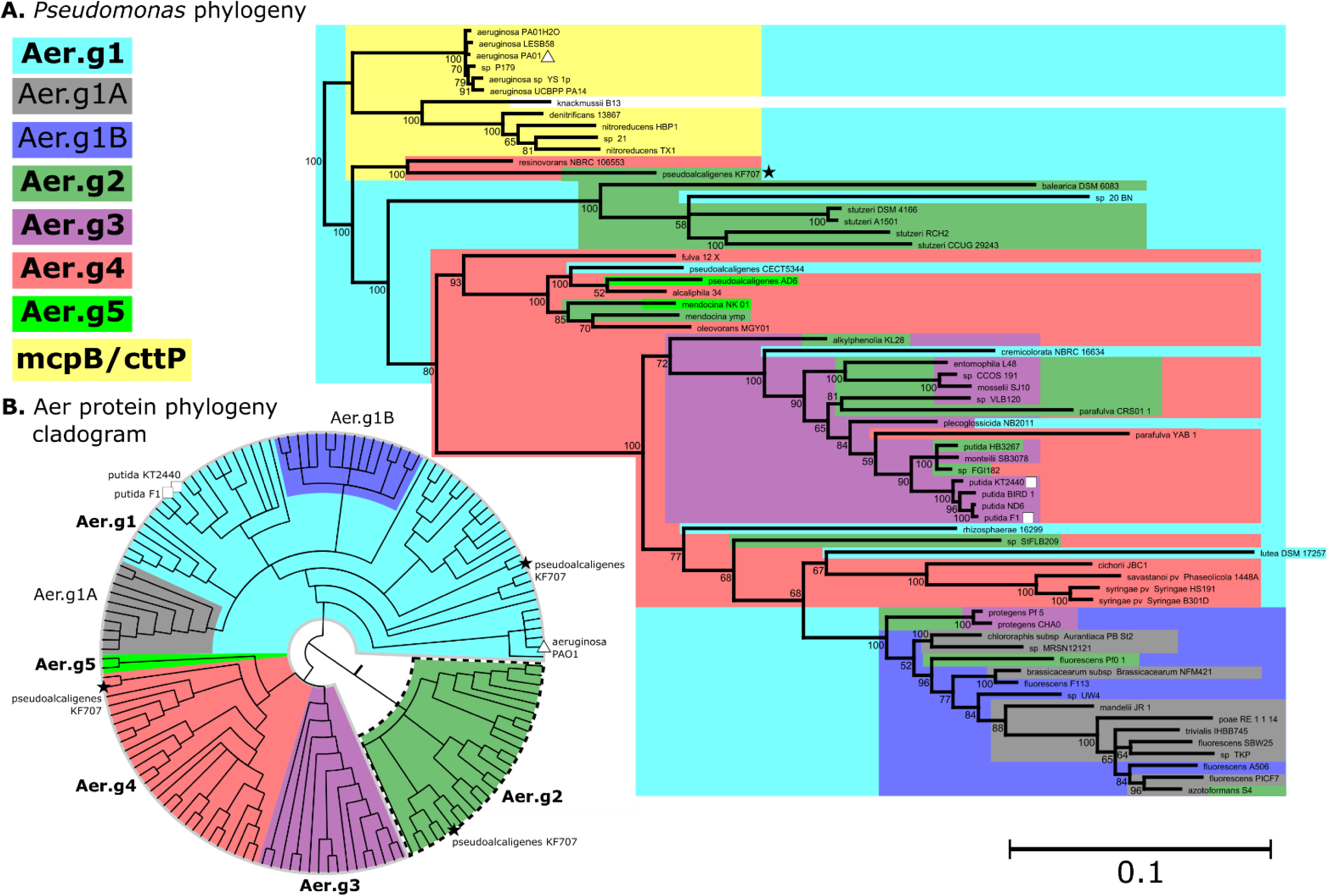
Phylogeny of select *Pseudomonas* species (A) and their Aer protein sequences (B). Species phylogeny is based on nucleotide sequences of concatenated *gyrB*, *rpoB* and *rpoD*. Numbers along branches indicate bootstrap support values from 100 replicates and scale bar indicates average number of nucleotide changes per position. Colours indicate the presence of particular Aer groups, matching the grouping defined by the cladogram (B), which was confirmed in subsequent analysis. The root for the Aer protein tree was estimated to be between Aer.g2 (dotted outline) and the other groups based on inspection of unrooted phylogenies including Aer from *E. coli* or similar sequences from closely related non-*Pseudomonas* species. See Supplementary Figures 1-3 for details. For both trees, nodes with support values below 50 were collapsed. Previously characterized Aer sequences from *P. aeruginosa* (triangle) and *P. putida* (squares) are indicated, as well as those from *P. pseudoalcaligenes* KF707 (black stars).

### Distribution of Groups within *Pseudomonas*

To determine the prevalence of each Aer homolog group within the *Pseudomonas* genus, the number of homologs from each group that each species possessed was counted and a hierarchically clustered heatmap was generated (Supplementary Figure 6). This analysis showed that all included species, except *P. denitrificans* ATCC13867, had an Aer.g1 homolog. Conversely, only species related to *P. aeruginosa* had McpB and CttP, which were always found together. There were very few species that had McpB/CttP and multiple Aer homologs, only *Pseudomonas* sp 21, *P. nitroreducens* HBP1, *P. resinovorans* NBRC 106553 and *P. pseudoalcaligenes* KF707 fit this category. Aer.g1 was duplicated only in *P. fluorescens* and related subgroups 1-8 [17], and these duplications matched the prior subdivision of Aer.g1 into Aer.g1A and Aer.g1B. Duplications of Aer.g2 occurred only (and always) in *P. stuzeri* and the closely related *P. balearica*. Unlike the other duplications, these were tandem duplications. Duplications of Aer.g3 only occurred in *P. protegens* CHA0 and Pf-5.

### Relationship between *Pseudomonas* and Aer Phylogenies

To understand how the phylogeny of Aer related to the phylogeny of *Pseudomonas*, a species phylogeny was generated (Figure 1A) using concatenated alignments of *gyrB*, *rpoB*, and *rpoD*. This phylogeny matched well with previously published *Pseudomonas* phylogenies [17, 27]. Examination of how the various Aer subgroups, as well as McpB/CttP, mapped onto the genus phylogeny revealed their likely evolutionary trajectories (Figure 1A). The presence of *aer*.*g1* in all species, except *P. denitrificans* indicates that it is a basal *Pseudomonas* gene. Both subdivisions of Aer.g1 occurred only in the more derived *P. fluorescens* subgroup. Conversely, *aer.g2* is distributed throughout the genus with no clear pattern. It is present in the more ancestral *P. stutzeri* group as well as the more derived *P. fluorescens* subgroup but is not present in many species in between. *aer*.g3 is present, except for *P. protegens*, exclusively in the *P. putida* group, though not all members of the clade have it. *aer*.g4 is present mostly in the related *P. straminea, P. oleovorans, P. putida, P. lutea* and *P. syringae* subgroups. Notably, it is not present in the *P. chlororaphis* or *P. fluorescens* subgroups, despite these being more derived than the aforementioned groups. *aer.g4* was also found in *P. resinovorans* and *P. pseudoalcaligenes* KF707, but not the rest of their *P. aeruginosa* subgroup. *aer.g5* is only present in two species, both in the *P. oleovorans* subgroup. *mcpB* and *cttP*, which were always found together, were exclusively present in the *P. aeruginosa* subgroup.

### Genetic Features of *aer* Homologs

The grouping of Aer into 5 subgroups based on protein sequence information was confirmed and expanded on by examining features of the underlying *aer* genes. The upstream and downstream regions of the *aer* genes were inspected to determine whether the genomic context was consistent across species (Figure 2). Additionally, the first two upstream and downstream genes of each *aer* gene were identified (Supplementary Table 1). The frequency of each of the associated genes and the general genomic context for each Aer homolog were thus examined to identify the most commonly associated genes (summarized in Figure 2). Each homolog was part of an apparently unique gene cluster: Aer.g1 with an aconitate hydratase (in 69% of species) and a CAAX amino protease (33%), Aer.g2 with a PAS-containing diguanylate cyclase/phosphodiesterase (88%), Aer.g3 with a different PAS-containing diguanylate cyclase/phosphodiesterase (88%) and Aer.g4 with a LysR-type transcriptional regulator (64%) and a DTW-domain containing protein (74%). Beyond these, the second upstream and downstream genes tended to vary more, though some upstream genes were clearly conserved: Aer.g1, 23S rRNA methyl transferase (42%); Aer.g3, C4 dicarboxylate transporter (63%); Aer.g4, agmatinase (56%).

**Figure 2:**
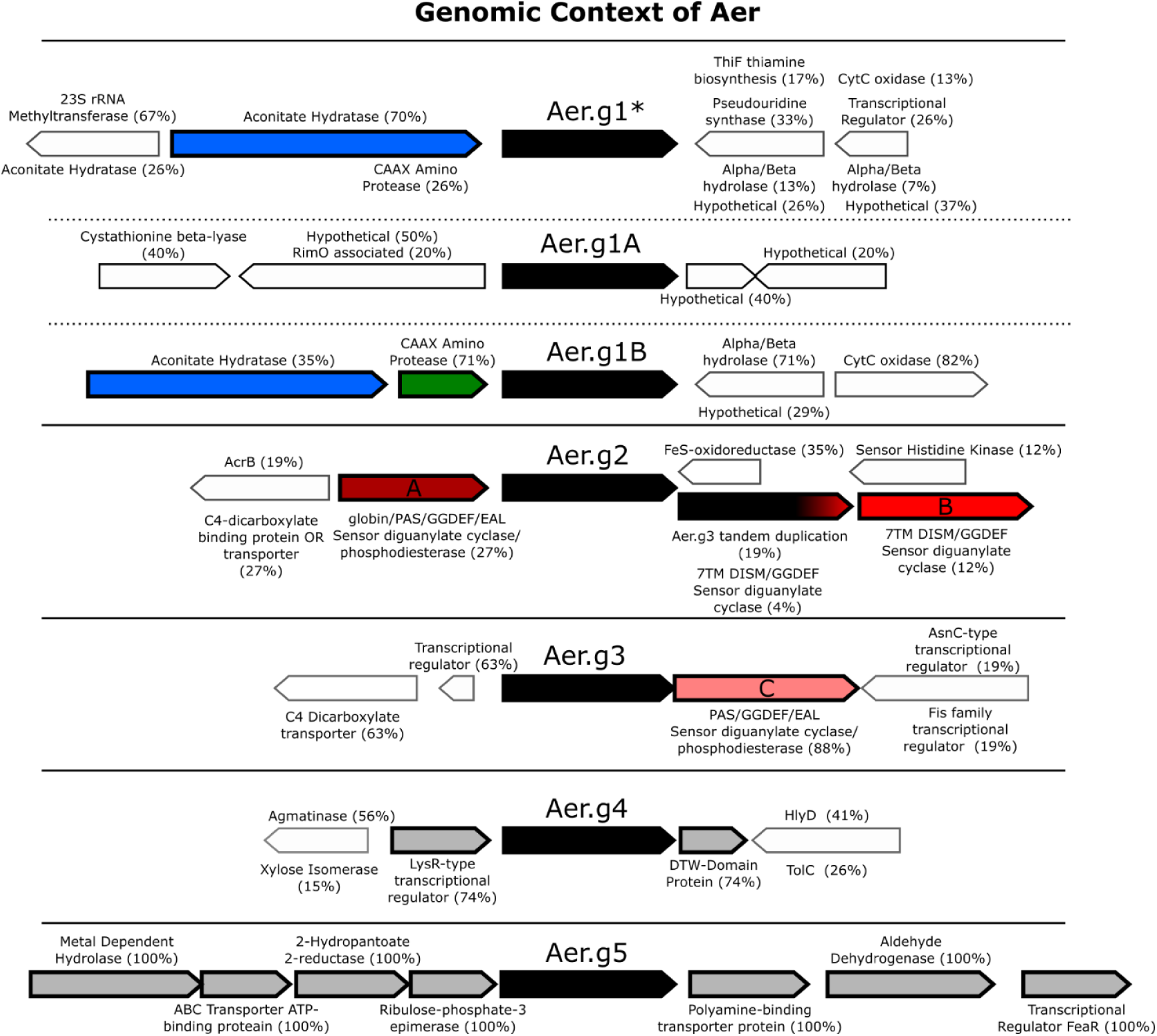
Frequency of occurrence and orientation of genes upstream and downstream from *aer* homologs of select *Pseudomonas* species. Genes with the same orientation and high frequency of occurrence with *aer* are outlined. Genes with noteworthy functions are coloured, and are discussed in the text, such as the 3 different diguanylate cyclases/phosphodiesterases (A, dark red; B, red; C, light red). Numbers beside gene functions indicate the frequency that they were found within each group or sub group. Gene lengths are approximate, *aer* is about 1.5kb long. Aer.g1* indicates Aer.g1 sequences not including Aer.g1A and Aer.g1B.

### Evidence for Horizontal Transfer of *aer* Homologs

While examining the genomic contexts of *aer* homologs, indicators of mobile genetic elements were found nearby in many cases (Supplementary Table 2). Along with the varied distribution of Aer homologs within the *Pseudomonas* genus, and the findings from the genus phylogeny that distantly related species had similar complements of *aer* genes, this implied that they may have been subject to horizontal transfer. Initial inspection of the Aer phylogeny identified 3 sequences that diverged from the expected species phylogeny. These 3 sequences (WP_015271024.1, *P. putida* HB3267; WP_014754514.1, *P. putida* ND6; and WP_013791017.1, *P. fulva* 12-X) were each found clustered with sequences from unrelated species whereas other Aer sequences were consistently clustered with sequences from closely related species. Of these 3 genes only the *aer.g3* gene from *P. putida* ND6 was associated with mobile elements. Mobile elements were found within 5 Kbp of *aer* homolog genes (Supplementary Table 2). When mobile elements were found, they were often located immediately up and/or downstream of the *aer* homolog and its associated gene(s) described in the above section. Tanglegrams were used to identify instances where the *gyrB*/*rpoB*/*rpoD* based species phylogeny did not match with the *aer* subgroup phylogenies (Supplementary Figures 7-10). Three instances for Aer.g2 and one for Aer.g3 were identified where the *aer* gene had likely been transferred horizontally. Probable instances of horizontal transfer were also identified from mapping the *aer* subgroups onto the genus phylogeny (Figure 1A). *P. resinovorans* and *P. pseudoalcaligenes* KF707 have Aer.g4 despite being distantly related to the other strains that possess this homolog. Also, both *P. protegens* strains have Aer.g3 despite being more distantly related to the *P. putida* subgroup strains that only have this homolog.

### Amino Acid Sequence Comparison of Groups

As genomic analysis of the Aer subgroups confirmed the AA sequence-based groupings from the SHMR analysis, further sequence analysis was pursued to uncover sequence features that distinguished the various Aer groups. First, the domain architecture from all sequences were compared (Supplementary Figure 11) by submitting all sequences to SMART [28]. All sequences were similar to the expected domain architecture of Aer from *P. aeruginosa* PA01, consisting of an N-terminal Per/Arnt/Sim (PAS) domain, transmembrane helices and then the cytoplasmic kinase control (called MA for methyl-accepting in SMART) region which includes the CheW/CheA interface. HAMP domains between the transmembrane region and MA domain were identified almost exclusively in Aer.g2 sequences, likely due to the difficulty in identifying this domain [29]. To further identify regions of the protein that are specific to the subgroups, results from the SHMR analysis for each inter-group comparison were plotted against the entire length of the Aer protein (Figure 3A) and compared to the overall conservation at each AA position (Supplementary Figure 12). As expected, the CheW interface region of the MCP-signal domain and the PAS ligand binding domain were the most conserved regions, and did not differ between groups. Inspection of the SHMR scores for each intergroup comparison showed that the regions determining the specificity of each group were the same for each Aer group (Figure 3A), except for Aer.g2, which had a group-specific region immediately N-terminal to the PAS domain. Sequences from this group also had variable length C-terminal extensions. The most notable group-specific region was in between the transmembrane helices and the beginning of the kinase control (MA) domain. As this is the location of the HAMP domain in *E. coli* Aer, this region was examined more closely.

**Figure 3:**
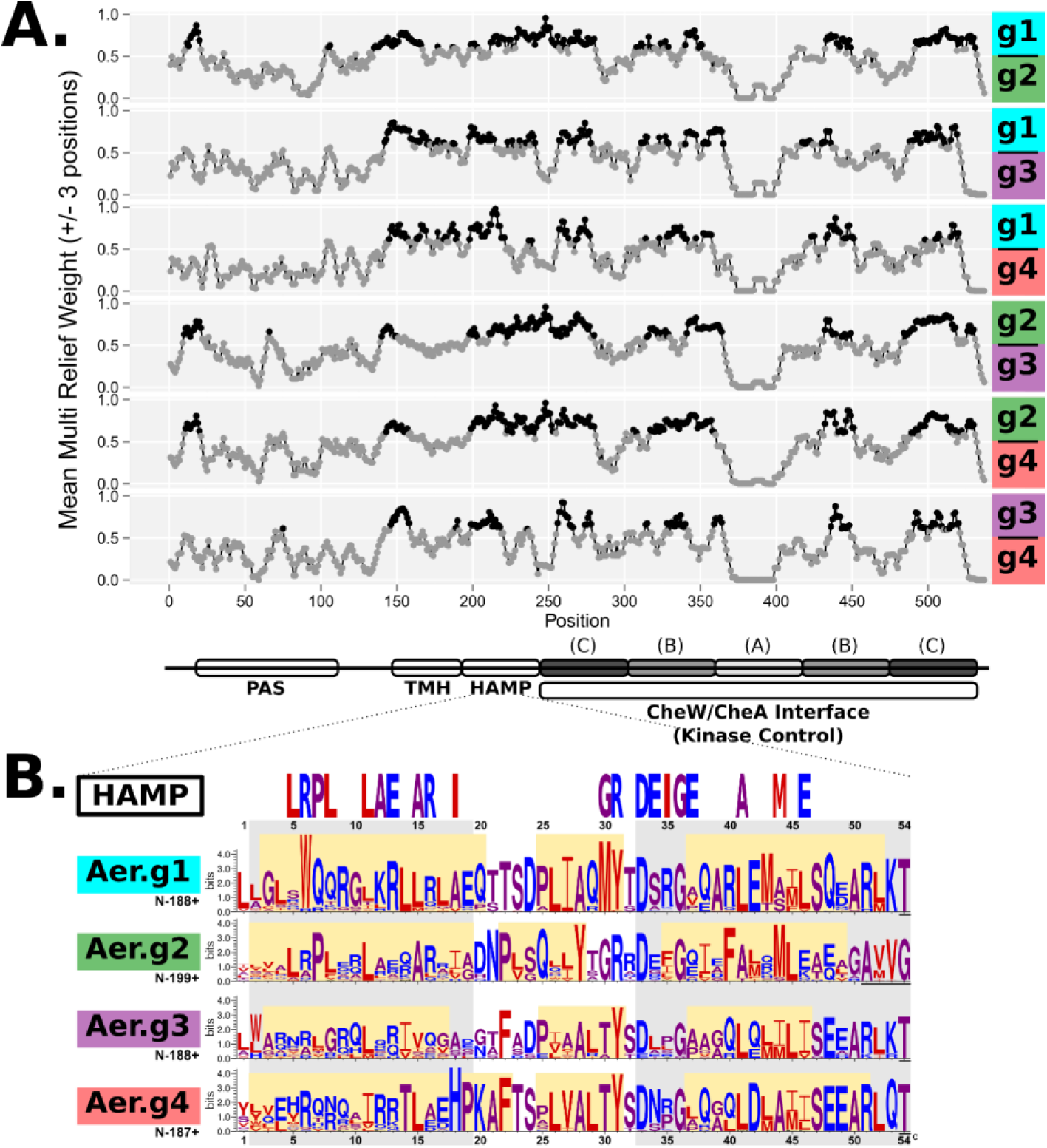
Multi-relief comparison of Aer groups showing group-specificity (A) and group-specific Weblogos of HAMP domains (B). Black points indicate positions with a multi-relief score above 0.6. This cutoff indicates positions that are conserved within groups but divergent between. Domains are PAS (Pern/Art/Sin) ligand binding, TMH (transmembrane helices), HAMP (Histidine kinases, adenylate cyclases, methyl-accepting chemotaxis proteins and phosphatases), and CheW/CheA Interface also called Kinase Control (subdivided into A, signaling; B, flexible bundle; C, methylation). The HAMP domain weblogos in B were defined as the 54 AA anchored around Asp33. Characteristic HAMP domain residues were obtained from the SMART database, helices are marked in grey as defined in [1], and for each subgroup in yellow as determined by PSIPRED [2, 3]. Amino acids are coloured for hydrophobicity.

### Group-Specific Domain Differences

First, the CheW/CheA interface region was examined more closely as its subdomains have been clearly defined [30]. In bacterial chemoreceptors this domain consists of pairs of heptads, separated into three subdomains. Central to the domain is a glutamate residue which denotes the ‘zero’ position from which paired heptads are counted outwards until the C-terminus on one side and the end of the domain on the other. As there were 20 heptads to the C-terminus in all sequences, except Aer.g2 which had variable length extensions, all Aer from *Pseudomonas* fall in the 40H class [30]. The three sub-domains (from the center outwards), signaling, flexible bundle, and methylation were marked and used to compare the conservation (Supplementary Figure 13) and group-specificity (Supplementary Figure 14) of the various Aer groups. Conservation showed that corresponding heptads of the more central methylation subdomain were better conserved than the flexible bundle subdomain. This corresponded with indications of group-specific features in the flexible bundle subdomain and in the outer heptads (furthest from the signaling subdomain) of the methylation subdomain.

The PAS and HAMP domains were also compared between all groups. The PAS domains from all Aer groups were not notably different (Supplementary Figure 14). Conversely, the region where a HAMP domain was expected, but only identified by SMART in Aer.g2, differed between each Aer group (Figure 3B). The predicted 54 AA HAMP domains could be anchored by a conserved aspartate and glycine [31], though the 51^st^ and 54^th^ residue of this domain marked the first AA of the kinase control domain of Aer.g2 and the other groups, respectively. Aer.g2 matched the SMART database HAMP domain residues the closest. Though all groups differed substantially from one another in this region, secondary structure prediction [32, 33] identified helices in each group which matched the expected HAMP domain architecture [31]. In Aer.g1, Aer.g3 and Aer.g4, the final seven AAs of the domain were well conserved, forming a heptad with the motif SZEARL(K/Q). Only the Z was present in Aer.g2. With regards to the subdivision of Aer.g1, both the PAS and HAMP domains supported keeping all sequences as a single group (Supplementary Figures 15 and 16). Aer.g2 also differed from the other three groups at its C-terminus as the final heptad of the kinase control domain differed and its members feature an up to 9 AA extension (Supplementary Figure 17).

## Phenotypic Results

To investigate the functions of Aer homologs, CttP and McpB, and to see whether they were linked, deletion mutants were constructed in *P. pseudoalcaligenes* KF707. Single and combination deletions were constructed using 2-step homologous recombination, the suicide vector pG19II and SacB sucrose counter-selection [34]. To test energy-taxis, soft agar swim plates were used [35]. In this assay cells consume carbon source from a central inoculation point then expand outwards via a combination of division, chemotaxis and energy-taxis. Non-chemotactic strains cannot expand beyond the point of inoculation but non-energy tactic cells will produce a ring of expanding cells that is smaller than the wild-type (Supplementary Figure 18). To ensure that energy-taxis was being compared between the different mutants, the results were compared to a taxis-negative CheA::KmR insertional inactivation mutant [36], and the chemotactic ability of all strains was also confirmed in classical chemotaxis swim plates (Supplementary Figure 19). Comparison of these metabolism independent results to the metabolism dependent results of the energy-taxis swim plates allowed for differences in energy-taxis capabilities to be observed between strains. All 24 mutants generated in this study, along with the wild-type and CheA control were tested in triplicate in media containing pyruvate (Figure 4) or succinate (Supplementary Figure 20). Ring diameters were measured from the furthest point reached from the inoculation centre (Supplementary Figure 21), normalized to the wild-type, then tested for differences using Tukey’s Honest Significant Differences test (Supplementary Table 3).

**Figure 4:**
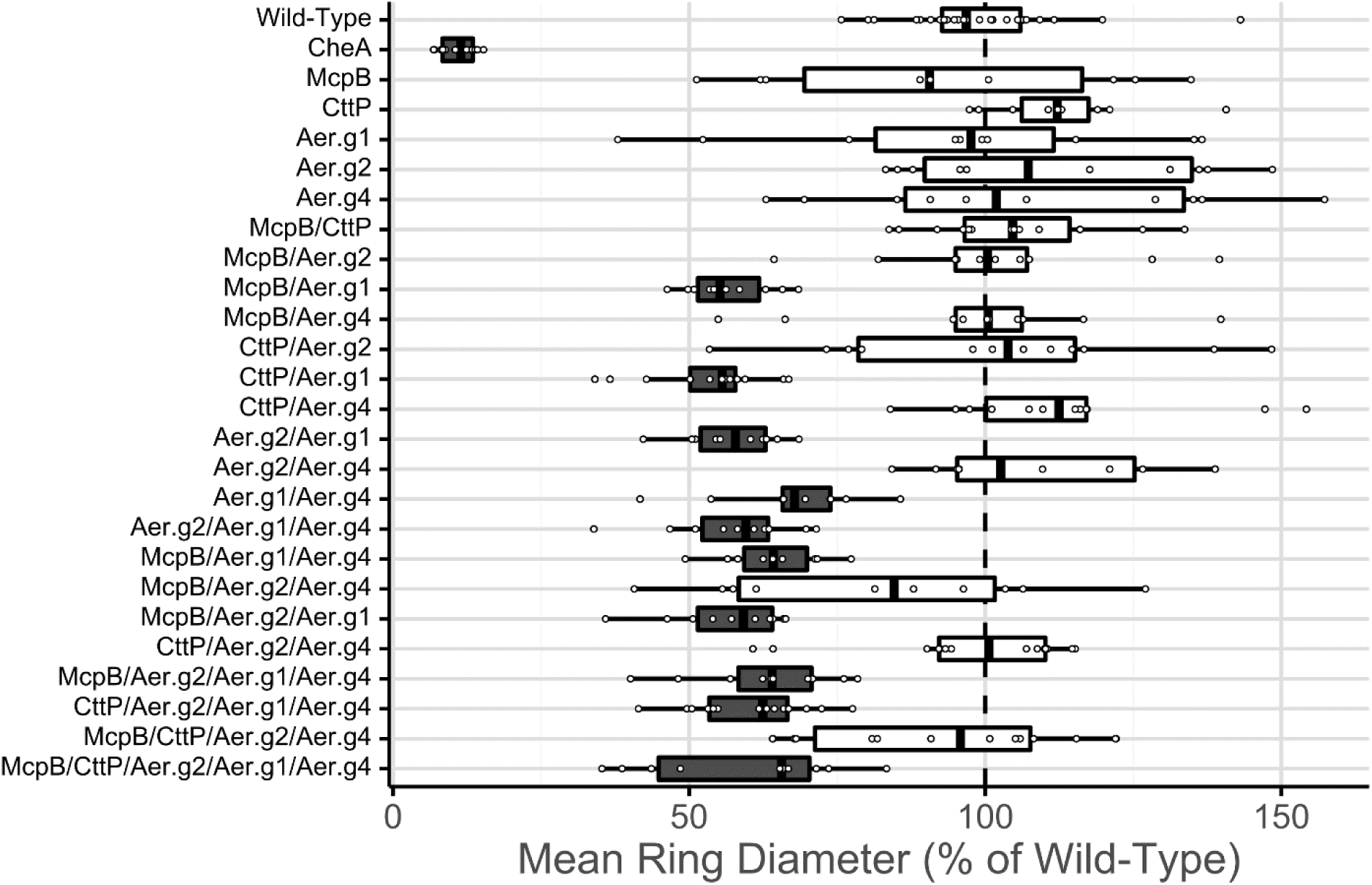
Mean normalized energy-taxis diameters in 50mM pyruvate of strains of *P. pseudoalcaligenes* KF707 with deletions of *mcpB, cttP* and *aer* homologs. Boxes filled in dark-grey indicate significant differences from the wild-type based on Tukey’s Honest Significant Differences test with a confidence value of 0.95. Values were normalized to the wild-type within each experiment at 24 and 48 h, allowing datapoints from both times to be combined (n ≥ 5). Wild-type strains were normalized to the mean of technical replicates within each experiment. The *cheA::KmR* mutant was not grown with antibiotic present.

The diameter of the chemotactic negative CheA mutant was about 10% of the wild-type, whereas energy-taxis negative mutants with diameters about 60% smaller were significantly different from the wild-type (Figure 4). All the singular deletions of Aer.g1, Aer.g2, Aer.g4, McpB and CttP had no effect on energy-taxis; only strains that had Aer.g1 deleted in combination with at least one other receptor showed changes in their energy-taxis phenotype. Conversely, deletion of any combination of these receptors, including the quadruple McpB/CttP/Aer.g2/Aer.g4 mutant did not adversely affect energy-taxis. The McpB/Aer.g2/Aer.g4 triple mutant had a diminished diameter, about 80% of the wild-type, but this was not significant. Results from plates containing succinate (Supplementary Figure 20) were similar to those obtained using pyruvate, but were more variable. The observed differences in swim diameters were not due to differences in growth rate or swimming speed as the changes in swim diameters between the 24 and 48 h measurements were similar between all strains (Supplementary Figure 22, Supplementary Table 4). Growth rates in liquid media also did not differ (Supplementary Figure 23). Additionally, all strains were tested for chemotaxis towards pyruvate, glucose (Supplementary Figure 19) and succinate (data not shown) using traditional swim plate assays. All strains except the CheA::KmR mutant were able to swim towards pyruvate, succinate and glucose. As Aer.g1 played a pivotal role in energy-taxis, the wild-type and quintuple McpB/CttP/Aer.g1/Aer.g3/Aer.g4 mutant were complemented with Aer.g1 on a plasmid which showed the expected increase and recovery, respectively, of energy-taxis in succinate plates (Supplementary Figure 24, Supplementary Table 5). Together these experiments showed that it was only the combined deletion of Aer.g1 with any of the other tested receptors that caused a reduction of energy-taxis.

## Discussion

### Protein sequence and genetics classify Aer homologs into 5 groups

Aer has previously been described in *P. putida* and *P. aeruginosa* [8, 37–39], though the two species differed in the number of apparent homologs indicating there is interspecies variation. Here we constructed a protein sequence-based phylogeny which separated Aer into 5 different groups. All groups varied in the same regions of the protein, except Aer.g2 which had a unique region at the C terminus, which does not resemble the C-terminal CheR interface pentapeptide of McpB [40]. The most conserved regions included the PAS ligand binding domain and CheW/CheA interface region which confer the necessary functions of FAD cofactor binding/signal reception [41] and signal transduction, respectively. The conserved regions in the signaling region were better conserved than the predicted PAS domain, though these domains are known to differ at the sequence level [41]. Interestingly, it was the cytoplasmic region between the transmembrane helices and kinase control domain where a HAMP domain was expected that differed most between groups (Figure 3). In Aer from *E. coli* the HAMP domain in this region [42] interacts with the PAS domain and is essential for the function of the protein as the HAMP domain transfers the signal through the protein [11]. The motif (5)RP[…]DEXG(36) is representative of ‘canonical’ HAMP domains where the signal is transduced through the membrane [31], but was only found in Aer.g2. As the signal in Aer can be passed directly from the cytoplasmic PAS domain to the HAMP domain as in *E. coli* [43], the HAMP domains of the other Aer groups were likely able to evolve without disrupting this function, as predicted [31]. This implies that Aer.g1, Aer.g3, Aer.g4 and Aer.g5 have diverged from their common ancestor with Aer.g2, which matches the phylogenetic results.

### Aer.g1 is the most prevalent homolog

Aer from *E. coli* has been well-studied, so was used to estimate the root of the *Pseudomonas* Aer phylogeny. After adding its sequence to the *Pseudomonas* alignment and removing any gaps created (*E. coli* Aer is of the 36H class), the subsequent phylogeny placed it in between Aer.g1 and Aer.g2 (Supplementary Figure 2). To confirm that this was an appropriate branch to root the tree, an additional phylogeny was generated adding closely related non-*Pseudomonas aer*-like genes (Supplementary Figure 3). Again, most sequences placed between Aer.g1 and Aer.g2 indicating that the Aer.g1/3/4/5 and Aer.g2 groups diverged from a common ancestor.

Aer.g1 was the most prevalent as all the *Pseudomonas* species included in this study had at least one homolog except *P. denitrificans* ATCC13867. As this species did not have any Aer homologs, this means that Aer is not part of the *Pseudomonas* core genome, though it is likely ancestral and was lost in *P. denitrificans*. Unlike Aer.g1, Aer.g2 was not present in all *Pseudomonas* species, but it was present in species from most subgroups. This suggests that both Aer.g1 and Aer.g2 were present in the last common ancestor of all *Pseudomonads*.

Almost all *aer.g1* genes were adjacent to an aconitase gene (Figure 3), indicating that they may be co-regulated or co-transcribed. In *Helicobacter pylori*, TlpD, which controls tactic behaviour in low-energy conditions, has been found to interact with aconitase [44], possibly indicating some common connection between aconitase and energy-taxis receptors. A gene encoding a predicted CAAX amino protease was found adjacent to one third of Aer.g1 homologs. This type of membrane-bound protease aids in prenylating proteins to ease their membrane localization [45]. The Aer.g1 proteins whose genes were located beside these predicted proteases did not have the expected CAAX motif and there were no obvious features at the sequence level that distinguished these sequences from other Aer homologs indicating that the genomically co-localized CAAX protease may not be involved in their functioning.

### Other Aer groups may have alternate functions

As the 5 previously characterized Aer receptors are Aer.g1 homologs with the same energy-taxis function, it is possible that the other homolog groups may have a related, but different and/or more specific function. The three Aer homologs of *P. putida* KT2440 were found to be differentially expressed [10], supporting this hypothesis. In *E. coli* Aer is also a thermosensor [46] indicating another possible role, or they could aid in tuning the energy-taxis response similarly to how Tsr and Tar from *E. coli* enable taxis towards the ideal pH from both higher and lower initial pHs [47]. The function of the additional Aer homologs could also be related to biofilm formation as Aer.g2 and Aer.g3 were each associated with their own PAS-domain containing diguanylate cyclase/phosphodiesterase. Though Aer is expected to only act as a Che1 (F6) chemosensor [48], the recent revelation that not all chemosensors mediate chemotaxis [49], and are instead involved in so-called ‘alternative cellular functions’ such as cyclic di-GMP signaling, makes this frequent association of phosphodiesterase/diguanylate cyclases with Aer.g2 and Aer.g3 noteworthy. Deletion of the diguanylate cyclase/phosphodiesterase associated with Aer.g3 in *P. putida* KT2440 caused a general defect in motility [10] and was later shown to be a bi-functional cyclic di-GMP phosphodiesterase and diguanylate cyclase [50]. These two homolog groups may thus be involved in energy-sensing behaviour in biofilms as c-di-GMP is an important regulator in the transition from planktonic to sessile growth modes. In *P. aeruginosa*, increased levels of cyclic di-GMP decreases the frequency of flagellar motor switching through MapZ as it inhibits CheR1 from methylating some MCPs [51]. As this shows a direct connection between chemotaxis and cyclic di-GMP, understanding the function of the Aer homolog-associated cyclase/phosphodiesterases will be of interest.

### Possibility of horizontal transfer of *aer* homologs

Chemoreceptors have been posited to be subject to horizontal gene transfer [52], as has been demonstrated in plasmids of *E. coli/Shigella* [53]. Here we presented multiple lines of evidence that suggest genes from the *aer* group have been horizontally transferred in the chromosome of *Pseudomonas* species. In the Aer phylogeny, three homologs from *P. putida* HB3267, ND6 and *P. fulva* 12-X each clustered beside Aer sequences from unrelated species indicating they may have been obtained from those species. Incongruencies between the Aer phylogeny and genus phylogeny were also found using tanglegrams. These figures highlight other instances where *aer* homolog sequences were more closely related than the species hosting these genes, which could indicate non-vertical inheritance. Additionally, we noted several cases where strains of the same species differed in which Aer homologs they possessed.

The observed variation in the distribution of Aer homologs is unsurprising as even within the *P. aeruginosa* pan-genome from 7 strains there are 2,000 accessory genes compared to 5,000 core genes [54]. The fact that *aer.g1* almost qualifies as part of the genus core genome (*P. dentrificans* being the only outlier) indicates that is a very useful gene in the varied environments inhabited by *Pseudomonas*. The existence of the various homologs speaks to their utility as they each may confer a more specific function. As many *Pseudomonas* species live in the soil and rhizosphere, which promotes horizontal gene transfer [55], there would be ample opportunity for these *aer* homolog genes to spread. Interestingly they appear to be restricted to *Pseudomonas* as reverse BLAST searches consistently returned *Pseudomonas* sequences as the closest hits.

### Genes influencing energy-taxis in *P. pseudoalcaligenes* KF707

*P. pseudoalcaligenes* KF707 was the only species that had three Aer homologs as well as McpB and CttP. We have previously examined its chemotactic ability towards biphenyl [56], and how its multiple *cheA* genes affect biofilm formation [36] and swimming motility [18]. Aer from *P. aeruginosa* PA01 [8], *P. putida* F1 [15] and *P. putida* KT2440 [10] have been previously characterized but are all part of the Aer.g1 homolog group. In the present study, all three *aer* homologs in *P. pseudoalcaligenes* KF707, representing *aer.g1*, *aer.g2* and *aer.g4*, as well as *mcpB* and *cttP* were deleted individually and in combination. The energy-taxis phenotypes of these mutants were compared based on their diameters in soft agar energy-taxis plates. As the growth rate of all the strains in these plates, and in liquid culture did not differ, as well as their chemotactic ability in swim plates, the observed differences were taken as indication that the perturbed genes influenced energy-taxis in some way.

No single deletions affected the energy-tactic ability of *P. pseudoalcaligenes* KF707. This differs from *P. putida* KT2440 which became energy-taxis negative when only its Aer.g1 homolog was inactivated [38]. Both species have Aer.g4, but KF707 has Aer.g2 and KT2440 Aer.g3. In KF707, co-deletion of either *aer.g4* or *aer.g2* in the Δ*aer.g1* background caused a reduction of energy-taxis, but deletion of *aer.g2* and *aer.g4* together had no effect. The three Aer homologs in KT2440 are differentially expressed and thus were not truly redundant [38], but in KF707 its Aer homologs appear to be at least partially redundant. In *Ralstonia solanacearum* both of its “Aer1” and “Aer2” receptors were necessary for aerotaxis and restored energy-taxis in a deficient *E. coli* mutant [57]. As these receptors are similar to Aer from *Pseudomonas* and *E. coli*, it was likely energy-taxis enabling the swimming of *R. solanacearum* towards oxygen. Interestingly these two receptors appear not to be redundant as inactivation of either one caused a defect. In *Vibrio cholerae*, the number of Aer homologs also differs between species. The El tor biotype has three, compared to two in strain O395N1 [58]. The additional Aer in the El tor biotype is most similar to Aer.g2, though all the sequences from *Vibrio* and *Ralstonia* appear to be quite different (identity below 50% on <90% coverage) than *Pseudomonas* sequences making it difficult to categorize these homologs according to the naming scheme developed here for *Pseudomonas.* Only one homolog in *V. cholera* was found to affect energy-taxis, leaving the exact function of the other homologs unknown, similar to *Pseudomonas*.

Aer and McpB were first characterized as aerotaxis receptors in *P. aeruginosa* PA01 [8]. McpB has been shown to bind oxygen [9], and its overexpression in *P. aeruginosa* and *E. coli* abolished chemotaxis [59] but its function as an aerotaxis receptor remains inconclusive. Here we showed that single deletion of *mcpB* did not affect energy-taxis, but co-deletion with *aer.g1* resulted in a similar reduction in energy-taxis as co-deletion with either of the other *aer* homologs. The McpB protein has a C-terminal extension that specifically allows CheR2, and only CheR2, to methylate it [40], and is required for the complexation of other Che2 proteins [13], making it appear that the entire Che2 pathway exists to transduce the signal sensed by McpB. This was recently confirmed using sequence analysis of all *P. aeruginosa* PAO1 chemoreceptors [48]. Our results were thus surprising as no connection was expected between McpB and the che1 system, which transduces the signal of Aer, though the functions of Aer and McpB in aerotaxis were linked in their initial characterization [8]. As their phenotypic outputs may overlap, *P. denitrificans* may be an ideal organism to study McpB, as it has no Aer homologs.

The *che2* gene cluster is preceded by *cttP* so we investigated the possibility of its involvement in energy-taxis as well. In *P. aeruginosa* PA01, CttP was characterized as a receptor for positive chemotaxis towards tetra/tri-chloroethylene (PCE/TCE) [19]. As this organism does not perform reductive dehalogenation, intentional taxis towards these compounds is puzzling. In *E. coli*, the receptors Tar and Tsr can mediate attractant and repellent responses to phenol through its direct interaction with the transmembrane helices or cytoplasmic HAMP signaling domains [20]. We thus hypothesized that a similar interaction may have mediated the observed PCE/TCE chemotaxis and that CttP actually has a different function. CttP is only found in organisms with McpB, and the gene is located immediately upstream of the *che2* gene cluster. Here we showed that co-deletion of *cttP* and a*er.g1* in *P. pseudoalcaligenes* KF707 reduced energy-taxis comparable to co-deletion of *aer.g2* or *aer.g4*. CttP has an unusual domain architecture compared to other MCPs as it has no predicted ligand-binding domain and a single N-terminal predicted transmembrane domain [60]. It does have the expected CheW/CheA interface domain, but also a long (>100 AA) C-terminal extension not seen in most MCPs. It could be possible that CttP does not directly sense any signal, but interacts with MCP receptor complexes and/or influences their interaction with the Che complex. Overexpression of *cttP* abolished all chemotaxis in both *P. aeruginosa* and *E. coli* [59], possibly indicating that its function is in modulating signaling from the che1 system.

Despite being the first gene in the *che2* gene cluster, *cttP* is involved only with *che1* genes as the *che2* system is dedicated to processing the signal sensed by *mcpB* [48]. It is thus puzzling that *cttP* was always found in species that have *mcpB*, and that deletion of *mcpB* in combination with *aer.g1* diminished energy-taxis in *P. pseudoalcaligenes* KF707. This could be due to cross-talk between the systems, which is supported by a recent study that showed that the *che2* system is an ancestral F7 chemosensory system that evolved and co-opted parts of the *che* (F6) system to become the well characterized system present in *E. coli* [61]. Here it was suggested that since the F7 system appears to function in stress response, but that CttP co-evolved with it and interacts with the F6 system[13] it may bridge the input/output of the two systems.

## Conclusions

In *Pseudomonas*, Aer is not a single receptor for energy-taxis, but a family of receptors which differ mostly in the HAMP domain. The number of Aer homologs varies widely between species and they have likely been horizontally transferred. The most prevalent homolog in the *Pseudomonas* genus, here named Aer.g1, mediates energy-taxis in *P. aeruginosa* and *P. putida*, but in *P. pseudoalcaligenes* KF707 Aer.g2 and Aer.g4, can also influence energy-taxis. McpB and CttP also appear to have some effect. These findings indicate that Aer and energy-taxis in *Pseudomonas* are more variable and complicated than the single receptor found in the archetypal species *P. aeruginosa*.

## Materials and Methods

### Protein Sequences

All sequences were obtained using NCBI databases and tools [62]. BLAST searches on the NCBI website [21] (pBLAST for draft and completed genomes, tBLASTn for whole-genome shotgun genomes) were performed using *P. aeruginosa* PAO1 Aer (NP_0250252.1) as a query sequence. From each species, all hits with >95% sequence coverage, no matter how low the sequence identity, were selected for inclusion. BLAST did not return any results with coverage values between 67% and 95%, indicating that all included sequences were likely to truly be Aer sequences. Expect values were always below 1×10^-100^. Four sequences were removed as they were redundant entries resulting from incorrect start site annotations resulting in two proteins with the same C-terminus but slightly different N-termini. Sequence accession numbers were then obtained from the international nucleotide sequence database collaboration (INSDC) [63].

### Amino Acid Alignment

Full length sequences were aligned using COBALT (on the NCBI website) [22] with default settings. Gaps were manually removed by adjusting the alignment in Jalview 2.10 [64] then re-aligning using MUSCLE with default settings [65]. Names were cleaned up using a custom script in R to reduce the names to just the species, strain and accession number in a presentable fashion [66]. Aer from *E. coli* (AGX35053.1), and seventeen sequences obtained by BLAST query using PAO1 Aer (NP_250252.1) against *Chromohalobacter salexigens, Marinobacter hydrocarbonoclasticus VT8, Marinobacter sp. ELB17, Hahella chejuensis KCTC 2396, Marinomonas sp. MWYL1, Marinomonas sp. MED121, Neptuniibacter caesariensis* [*Oceanospirillum sp. MED92*]*, Bermanella marisrubri* [*Oceanobacter sp. RED65*]*, Reinekea blandensis (sp. MED297)* were added to the *Pseudomonas*-only alignment which was then re-aligned and all gaps trimmed so phylogenies were built only based on conserved regions. Maximum likelihood phylogenies were built using PHYML [23], as implemented by the South of France Bioinformatics platform [24]. PHYML options: Amino-Acids, 100 bootstrap replicates, JTT amino-acid substitution model, BEST tree topology search operation, tree topology, branch length and rate parameters were optimized. Consense was then used to generate a consensus tree from the bootstrap replicates using the majority rule (extended).

### Sequence Harmony and Multi-Relief Analysis to Compare Groups

The multi-Harmony server was used to apply sequence harmony and multi-relief (SHMR) to validate groupings made based on the ML tree grouping and alignment. Groups were manually decided based on the tree topology and corresponding alignment, initially making for 7 groups. These seven groups were compared in pair-wise fashion using SHMR [26]. Unique regions of each group of Aer homologs were identified by comparing the SHMR scores for each pair-wise comparison with the overall conservation score of each AA and the domain architecture of the Aer protein. Conservation scores were obtained from Jalview [64], and along with the SHMR scores were smoothed and plotted using ggplot2 [67] in R. Smoothing was performed by calculating the average of the 3 proceeding and following AAs for each position. The domain architecture of Aer was obtained from the conserved domain database using Aer from *P. aeruginosa* as a query (NP_250252.1) (CDD) [68]. WebLogos were generated using the Weblogo generator tool [69].

### Distribution of Groups

Based on the ML tree grouping, the number of times a strain appeared in each group was counted. A hierarchically clustered heatmap using Bray-Curtis distance and average clustering was made in R. Presence of Aer-2 (NP_248866.1) and CttP (WP_003106690.1) were determined using BLAST searches specifically against the strains.

### Detection of Evidence of Horizontal Gene Transfer

Graphical representations of the Aer homologs nucleotide sequences were manually inspected on NCBI. The first two genes upstream and downstream of the *aer* homolog were noted, along with any mobile elements (transposase, integrase and inverted repeats) within 5kb. For each different upstream and downstream gene their frequency of occurrence was calculated for each homolog group. The frequency of occurrence of each type of mobile element was also calculated, as well as for the complete set of homologs.

### Generation of DNA Sequence Phylogeny

Gene sequences were obtained by BLAST [21] using sequences of *gyrB, rpoB* and *rpoD* from *P. aeruginosa* PAO1. Sequences were aligned separately by MAFFT [70] and positions with gaps in any strain were removed. Sequences from all three genes were then concatenated and a ML phylogeny was generated using DIVEIN [71]. DIVEIN parameters: substitution model, GTR; equilibrium frequencies, optimized; proportion of invariable sites and gamma distribution parameter, estimated; number of substitution rate categories, 4; tree searching, NNI+SPR; tree optimized for topology and branch lengths; 100 bootstrap replicates. Tanglegrams were generated in Dendroscope [72] by pruning branches from the genus DNA tree so that only strains that matched the particular Aer homolog group were included.

### Culture Growth

For molecular biology, cultures were routinely cultured in lysogeny broth (LB, 5 g/L yeast extract, 10 g/L Tryptone, 10 g/L NaCl). For energy-taxis experiments *P. pseudoalcaligenes* KF707 strains were grown overnight (16 h) in minimal salts media containing 10 mM pyruvate or succinate. Minimal salts media contained (in g/L) K_2_HPO_4_, 3; NaH_2_PO_4_, 1.15; NH_4_Cl, 1; KCl, 0.15; MgSO_4_, 0.15; CaCl_2_, 0.01; FeSO_4_, 0.0025. The latter four were sterile filtered and added after autoclaving.

### Generation of Deletion Constructs and Mutants

Deletions were generated using two-step homologous recombination [34] with pG19II[73]. Nucleotide sequences for Aer.g1, Aer.g2, Aer.g4, McpB and CttP were obtained from the draft genome sequence of *P. pseudoalcaligenes* KF707 [74]. Genomic DNA was isolated by the phenol/chloroform method [75]. Primers were designed using Primer BLAST [76] and purchased from IDT (IDT, USA). Deletion constructs with BamHI or HinDIII restriction sites and complementary overlaps were ligated into pG19II using T4 ligase (Invitrogen, USA) and transformed into *E. coli* Top10F’ using standard methods [75]. Sequencing (Eurofins, USA) confirmed colonies were puddle mated with *P. pseudoalcaligenes* KF707 and *E. coli* HB101 carrying the helper plasmid pRK2013 [77]. Transconjugants were selected on AB glucose Gm20 plates. Second recombination events were selected for by pre-growth on LB no salt then selection on LB 10% sucrose. Colony PCR and sequencing (Eurofins, USA) were used to confirm the deletion. *P. pseudoalcaligenes* KF707 *cheA::Tn5* from Tremaroli *et al* 2011 [36] is called the ‘cheA’ mutant throughout the text as its cheA1 (equivalent to *P. aeruginosa* PA01) has been disrupted.

### Energy-taxis Swim Plates

Swim plates were made by making minimal salts media with 0.3% agar and 50mM succinate or pyruvate. Strains were grown overnight in MSM containing 10mM of the appropriate carbon source. Plates were inoculated by dipping a sterilized needle into the culture then stabbing into the plate. The diameter of growth for each strain was measured at the maximum distance away from the inoculation centre that bacteria were visible, at 24 and 48 h either manually using a ruler or digitally using a photograph and ImageJ [78]. Experiments were repeated at least 3 times for all strains in each media, always including at least one wild-type to be used as a normalizing control. Collected data were processed in R to normalize the size of the growth diameter to the corresponding wild-type size at 24 and 48 h. Tukey’s Honest Significant Differences test was used to determine if the differences between strains were significant for each carbon source. As the TukeyHSD() function in R [66] tests ALL pair-wise comparisons, only those comparing each strain to the wild-type are presented here. However, this makes these results more robust as the false positive correction for a confidence level of 0.95 was applied to all 325 comparisons which were tested.

### Chemotaxis Swim Plates

Strains were grown up overnight as before, then 1 mL was pelleted, washed once with 1 mL minimal salts media (no carbon source) then resuspended in 100 µL minimal salts media (no carbon source). 20 µL was spotted at the edge of a minimal salts media plate containing no carbon source and 0.3% agar. Either an agar plug containing 50 mM carbon source or small amount of crystals was placed in the centre and plates were incubated overnight at 30°C. Plates were photographed and positive chemotaxis was interpreted as an arc of cells nearer to the centre of the plate than the cells had been spotted.

### Growth Curves

Overnight cultures of all strains were normalized to OD 0.1 then diluted 1/100 into 200 µL minimal salts media with 10mM pyruvate in a 96 well microtiter plate. The plate was incubated at 30°C, shaking at 150RPM and the OD600 was checked every 6h.

### Complementation

Aer.g1 was complemented into the wild-type and the quintuple mutant (Aer-2/CttP/Aer.g1/Aer.g2/Aer.g4). Aer.g1 was cloned into pSEVA_342 [79], and was conjugated into *P. pseudoalcaligenes* KF707 the same as for the deletion constructs, only selection was performed on AB glucose plates with 30µg/mL chloramphenicol. For the energy-taxis assays, overnight cultures were grown with 30µg/mL chloramphenicol. Instead of directly comparing to the wild-type, these strains were compared based on their change in energy-taxis diameters over 24h. All plasmids used in this study are summarized in Supplementary Table 6.

## Acknowledgements

The authors would like to thank Stephana Cherak for her technical assistance, and Dr. Iain George and Dr. Gordon Chua for use of their imaging setup for photographing the swim plates. We thank Dr. Martina Cappelletti for assistance with the colony PCR design and her insights into KF707 physiology and genomic interpretation. This project was funded by Natural Sciences and Engineering Research Council (NSERC) of Canada Discovery Grant RGPIN/219895-2010 to RJT.

## References

1. Wadhams GH, Armitage JP. Making sense of it all: Bacterial chemotaxis. Nat Rev Microbiol 2004;5:1024–1037.

2. Sampedro I, Parales RE, Krell T, Hill JE. Pseudomonas chemotaxis. FEMS Microbiol Rev 2015;39:17–46.

3. Krell T. Tackling the bottleneck in bacterial signal transduction research: high-throughput identification of signal molecules. Mol Microbiol 2015;96:685–8.

4. Alexandre G, Greer-Phillips S, Zhulin IB. Ecological role of energy taxis in microorganisms. FEMS Microbiol Rev 2004;28:113–26.

5. Bibikov SI, Biran R, Rudd KE, Parkinson JS. A signal transducer for aerotaxis in Escherichia coli. J Bacteriol 1997;179:4075–4079.

6. Rebbapragada A, Johnson MS, Harding GP, Zuccarelli AJ, Fletcher HM, et al. The Aer protein and the serine chemoreceptor Tsr independently sense intracellular energy levels and transduce oxygen, redox, and energy signals for Escherichia coli behavior. Proc Natl Acad Sci U S A 1997;94:10541–6.

7. Samanta D, Widom J, Borbat PP, Freed JH, Crane BR. Bacterial Energy Sensor Aer Modulates the Activity of the Chemotaxis Kinase CheA Based on the Redox State of the Flavin Cofactor. J Biol Chem 2016;291:25809–25814.

8. Hong CS, Shitashiro M, Kuroda A, Ikeda T, Takiguchi N, et al. Chemotaxis proteins and transducers for aerotaxis in Pseudomonas aeruginosa. FEMS Microbiol Lett 2004;231:247–252.

9. Watts KJ, Taylor BL, Johnson MS. PAS/poly-HAMP signalling in Aer-2, a soluble haem-based sensor. Mol Microbiol 2011;79:686–99.

10. Sarand I, Österberg S, Holmqvist S, Holmfeldt P, Skärfstad E, et al. Metabolism-dependent taxis towards (methyl)phenols is coupled through the most abundant of three polar localized Aer-like proteins of Pseudomonas putida. Environ Microbiol 2008;10:1320–1334.

11. Garcia D, Watts KJ, Johnson MS, Taylor BL. Delineating PAS-HAMP interaction surfaces and signalling-associated changes in the aerotaxis receptor Aer. Mol Microbiol 2016;100:156–172.

12. Garcia D, Orillard E, Johnson MS, Watts KJ. Gas Sensing and Signaling in the PAS-Heme Domain of the Pseudomonas aeruginosa Aer2 Receptor. J Bacteriol 2017;199:e00003–17.

13. Güvener ZT, Tifrea DF, Harwood CS. Two different Pseudomonas aeruginosa chemosensory signal transduction complexes localize to cell poles and form and remould in stationary phase. Mol Microbiol 2006;61:106–18.

14. Nichols NN, Harwood CS. An aerotaxis transducer gene from Pseudomonas putida. FEMS Microbiol Lett 2000;182:177–183.

15. Luu R a., Schneider BJ, Ho CC, Nesteryuk V, Ngwesse SE, et al. Taxis of Pseudomonas putida F1 toward phenylacetic acid is mediated by the energy taxis receptor AER2. Appl Environ Microbiol 2013;79:2416–2423.

16. Bodilis J, Nsigue Meilo S, Cornelis P, De Vos P, Barray S. A long-branch attraction artifact reveals an adaptive radiation in pseudomonas. Mol Biol Evol 2011;28:2723–6.

17. Gomila M, Peña A, Mulet M, Lalucat J, García-Valdés E. Phylogenomics and systematics in Pseudomonas. Front Microbiol 2015;6:214.

18. Fedi S, Triscari-Barberi T, Nappi MR, Sandri F, Booth SC, et al. The Role of cheA Genes in Swarming and Swimming Motility ofPseudomonas pseudoalcaligenes KF707. Microbes Environ 2016;31:169–172.

19. Shitashiro M, Tanaka H, Hong CS, Kuroda A, Takiguchi N, et al. Identification of chemosensory proteins for trichloroethylene in Pseudomonas aeruginosa. J Biosci Bioeng 2005;99:396–402.

20. Pham HT, Parkinson JS. Phenol sensing by Escherichia coli chemoreceptors: a nonclassical mechanism. J Bacteriol 2011;193:6597–604.

21. Altschul SF, Gish W, Miller W, Myers EW, Lipman DJ. Basic local alignment search tool. J Mol Biol 1990;215:403–10.

22. Papadopoulos JS, Agarwala R. COBALT: Constraint-based alignment tool for multiple protein sequences. Bioinformatics 2007;23:1073–1079.

23. Guindon S, Gascuel O. A Simple, Fast, and Accurate Algorithm to Estimate Large Phylogenies by Maximum Likelihood. Syst Biol 2003;52:696–704.

24. Guindon S, Dufayard J, Lefort V. New Algorithms and Methods to Estimate Maximum-Likelihood Phylogenies : Assessing the Performance of PhyML 3.0. Syst Biol 2010;59:307–21.

25. Williams KP, Gillespie JJ, Sobral BWS, Nordberg EK, Snyder EE, et al. Phylogeny of gammaproteobacteria. J Bacteriol 2010;192:2305–14.

26. Brandt BW, Feenstra KA, Heringa J. Multi-Harmony: detecting functional specificity from sequence alignment. Nucleic Acids Res 2010;38:W35–40.

27. Özen AI, Ussery DW. Defining the Pseudomonas genus: where do we draw the line with Azotobacter? Microb Ecol 2012;63:239–48.

28. Schultz J, Copley RR, Doerks T, Ponting CP, Bork P. SMART: a web-based tool for the study of genetically mobile domains. Nucleic Acids Res 2000;28:231–4.

29. Airola M V., Watts KJ, Crane BR. Identifying divergent HAMP domains and poly-HAMP chains. Journal of Biological Chemistry. Epub ahead of print 2010. DOI: 10.1074/jbc.L109.075721.

30. Alexander RP, Zhulin IB. Evolutionary genomics reveals conserved structural determinants of signaling and adaptation in microbial chemoreceptors. Proc Natl Acad Sci U S A 2007;104:2885–90.

31. Dunin-Horkawicz S, Lupas AN. Comprehensive analysis of HAMP domains: Implications for transmembrane signal transduction. J Mol Biol 2010;397:1156–1174.

32. Zimmermann L, Stephens A, Nam SZ, Rau D, Kübler J, et al. A Completely Reimplemented MPI Bioinformatics Toolkit with a New HHpred Server at its Core. J Mol Biol. Epub ahead of print 2018. DOI: 10.1016/j.jmb.2017.12.007.

33. Jones DT. Protein secondary structure prediction based on position-specific scoring matrices. J Mol Biol. Epub ahead of print 1999. DOI: 10.1006/jmbi.1999.3091.

34. Hmelo LR, Borlee BR, Almblad H, Love ME, Randall TE, et al. Precision-engineering the Pseudomonas aeruginosa genome with two-step allelic exchange. Nat Protoc 2015;10:1820–1841.

35. Harwood CS, Nichols NN, Kim M, Diitty JL, Parales RE. Identification of the pcaRKF Gene Cluster from Pseudomonas. Biotechniques 1994;176:6479–6488.

36. Tremaroli V, Fedi S, Tamburini S, Viti C, Tatti E, et al. A histidine-kinase cheA gene of Pseudomonas pseudoalcaligens KF707 not only has a key role in chemotaxis but also affects biofilm formation and cell metabolism. Biofouling 2011;27:33–46.

37. Nichols NN, Harwood CS. An aerotaxis transducer gene from Pseudomonas putida. FEMS Microbiol Lett 2000;182:177–183.

38. Sarand I, Österberg S, Holmqvist S, Holmfeldt P, Skärfstad E, et al. Metabolism-dependent taxis towards (methyl)phenols is coupled through the most abundant of three polar localized Aer-like proteins of Pseudomonas putida. Environ Microbiol 2008;10:1320–1334.

39. Luu R a., Schneider BJ, Ho CC, Nesteryuk V, Ngwesse SE, et al. Taxis of Pseudomonas putida F1 toward phenylacetic acid is mediated by the energy taxis receptor AER2. Appl Environ Microbiol 2013;79:2416–2423.

40. García-Fontana C, Corral Lugo A, Krell T. Specificity of the CheR2 methyltransferase in Pseudomonas aeruginosa is directed by a C-terminal pentapeptide in the McpB chemoreceptor. Sci Signal 2014;7:ra34.

41. Henry JT, Crosson S. Ligand-Binding PAS Domains in a Genomic, Cellular, and Structural Context. Annu Rev Microbiol 2011;65:261–286.

42. Taylor BL. Aer on the inside looking out: paradigm for a PAS?HAMP role in sensing oxygen, redox and energy. Mol Microbiol 2007;65:1415–1424.

43. Watts KJ, Ma Q, Johnson MS, Taylor BL. Interactions between the PAS and HAMP Domains of the Escherichia coli Aerotaxis Receptor Aer. J Bacteriol 2004;186:7440–7449.

44. Behrens W, Schweinitzer T, McMurry JL, Loewen PC, Buettner FFR, et al. Localisation and protein-protein interactions of the Helicobacter pylori taxis sensor TlpD and their connection to metabolic functions. Sci Rep 2016;6:23582.

45. Pei J, Mitchell DA, Dixon JE, Grishin N V. Expansion of Type II CAAX Proteases Reveals Evolutionary Origin of γ-Secretase Subunit APH-1. 2011. Epub ahead of print 2011. DOI: 10.1016/j.jmb.2011.04.066.

46. Nishiyama S, Ohno S, Ohta N, Inoue Y, Fukuoka H, et al. Thermosensing function of the Escherichia coli redox sensor Aer. J Bacteriol 2010;192:1740–3.

47. Yang Y, Sourjik V. Opposite responses by different chemoreceptors set a tunable preference point in *Escherichia coli* pH taxis. Mol Microbiol 2012;86:1482–1489.

48. Ortega DR, Fleetwood AD, Krell T, Harwood CS, Jensen GJ, et al. Assigning chemoreceptors to chemosensory pathways in Pseudomonas aeruginosa. Proc Natl Acad Sci 2017;201708842.

49. Bardy SL, Briegel A, Rainville S, Krell T. Recent advances and future prospects in bacterial and archaeal locomotion and signal transduction. J Bacteriol 2017;199:e00203–17.

50. Österberg S, Åberg A, Herrera Seitz MK, Wolf-Watz M, Shingler V. Genetic dissection of a motility-associated c-di-GMP signalling protein of Pseudomonas putida. Environ Microbiol Rep 2013;5:556–65.

51. Xu L, Xin L, Zeng Y, Yam JKH, Ding Y, et al. A cyclic di-GMP–binding adaptor protein interacts with a chemotaxis methyltransferase to control flagellar motor switching. Sci Signal;9. http://stke.sciencemag.org.ezlibproxy1.ntu.edu.sg/content/9/450/ra102.full (2016, accessed 10 October 2017).

52. Zhulin IB. The superfamily of chemotaxis transducers: From physiology to genomics and back. Adv Microb Physiol 2001;45:157–198.

53. Borziak K, Fleetwood AD, Zhulin IB. Chemoreceptor gene loss and acquisition via horizontal gene transfer in Escherichia coli. J Bacteriol 2013;195:3596–602.

54. Qiu X, Kulasekara BR, Lory S. Role of Horizontal Gene Transfer in the Evolution of Pseudomonas aeruginosa Virulence. Genome Dyn 2009;6:126–39.

55. Sengeløv G, Kristensen KJ, Sørensen AH, Kroer N, Sørensen SJ. Effect of Genomic Location on Horizontal Transfer of a Recombinant Gene Cassette Between Pseudomonas Strains in the Rhizosphere and Spermosphere of Barley Seedlings. Curr Microbiol 2001;42:160–167.

56. Tremaroli V, Suzzi CV, Fedi S, Ceri H, Zannoni D, et al. Tolerance of Pseudomonas pseudoalcaligenes KF707 to metals, polychlorobiphenyls and chlorobenzoates: Effects on chemotaxis-, biofilm- and planktonic-grown cells. FEMS Microbiol Ecol 2010;74:291–301.

57. Yao J, Allen C. The plant pathogen Ralstonia solanacearum needs aerotaxis for normal biofilm formation and interactions with its tomato host. J Bacteriol 2007;189:6415–6424.

58. Boin MA, Häse CC. Characterization of Vibrio cholerae aerotaxis. FEMS Microbiol Lett 2007;276:193–201.

59. Ferrandez A, Hawkins AC, Summerfield DT, Harwood CS. Cluster II che Genes from Pseudomonas aeruginosa Are Required for an Optimal Chemotactic Response. J Bacteriol 2002;184:4374–4383.

60. Kim HE, Shitashiro M, Kuroda A, Takiguchi N, Ohtake H, et al. Identification and characterization of the chemotactic transducer in Pseudomonas aeruginosa PAO1 for positive chemotaxis to trichloroethylene. J Bacteriol 2006;188:6700–6702.

61. Ortega DR, Subramanian P, Mann P, Kjær A, Chen S, et al. Repurposing a macromolecular machine: Architecture and evolution of the F7 chemosensory system. bioRxiv 2019;653600.

62. Tatusova T, Ciufo S, Fedorov B, O’Neill K, Tolstoy I. RefSeq microbial genomes database: new representation and annotation strategy. Nucleic Acids Res 2014;42:D553–9.

63. Cochrane G, Karsch-Mizrachi I, Takagi T, International Nucleotide Sequence Database Collaboration IN. The International Nucleotide Sequence Database Collaboration. Nucleic Acids Res 2016;44:D48–50.

64. Waterhouse AM, Procter JB, Martin DMA, Clamp M, Barton GJ. Jalview Version 2--a multiple sequence alignment editor and analysis workbench. Bioinformatics 2009;25:1189–91.

65. Edgar RC. MUSCLE: multiple sequence alignment with high accuracy and high throughput. Nucleic Acids Res 2004;32:1792–1797.

66. R Core Team. R: A language and environment for statistical computing. R Found Stat Comput Vienna, Austria 2014;2014.

67. Wickham H. ggplot2. Wiley Interdiscip Rev Comput Stat 2011;3:180–185.

68. Marchler-Bauer A, Derbyshire MK, Gonzales NR, Lu S, Chitsaz F, et al. CDD: NCBI’s conserved domain database. Nucleic Acids Res 2015;43:D222–6.

69. Crooks GE, Hon G, Chandonia JM, Brenner SE. WebLogo: A sequence logo generator. Genome Res 2004;14:1188–1190.

70. Katoh K, Standley DM. MAFFT Multiple Sequence Alignment Software Version 7: Improvements in Performance and Usability. Mol Biol Evol 2013;30:772–780.

71. Deng W, Maust BS, Nickle DC, Learn GH, Liu Y, et al. DIVEIN: A web server to analyze phylogenies, sequence divergence, diversity, and informative sites. Biotechniques 2010;48:405–408.

72. Huson DH, Scornavacca C. Dendroscope 3: An Interactive Tool for Rooted Phylogenetic Trees and Networks. Syst Biol 2012;61:1061–1067.

73. Maseda H, Sawada I, Saito K, Uchiyama H, Nakae T, et al. Enhancement of the mexAB-oprM efflux pump expression by a quorum-sensing autoinducer and its cancellation by a regulator, MexT, of the mexEF-oprN efflux pump operon in Pseudomonas aeruginosa. Antimicrob Agents Chemother 2004;48:1320–8.

74. Triscari-Barberi T, Simone D, Calabrese FM, Attimonelli M, Hahn KR, et al. Genome sequence of the polychlorinated-biphenyl degrader pseudomonas pseudoalcaligenes KF707. J Bacteriol 2012;194:4426–4427.

75. Sambrook J, Fritsch EF, Maniatis T. No Title. Mol Biol A Lab Man.

76. Ye J, Coulouris G, Zaretskaya I, Cutcutache I, Rozen S, et al. Primer-BLAST: a tool to design target-specific primers for polymerase chain reaction. BMC Bioinformatics 2012;13:134.

77. Figurski DH, Helinski DR. Replication of an origin-containing derivative of plasmid RK2 dependent on a plasmid function provided in trans. Proc Natl Acad Sci U S A 1979;76:1648–52.

78. Schneider CA, Rasband WS, Eliceiri KW. NIH Image to ImageJ: 25 years of image analysis. Nat Methods 2012;9:671–675.

79. Silva-Rocha R, Martínez-García E, Calles B, Chavarría M, Arce-Rodríguez A, et al. The Standard European Vector Architecture (SEVA): A coherent platform for the analysis and deployment of complex prokaryotic phenotypes. Nucleic Acids Res 2013;41:D666–D675.

