## Supplementary Material for "Phylogenetic Characterization of the Energy-taxis Receptor Aer in *Pseudomonas* and Phenotypic Characterization in *P. pseudoalcaligenes* KF707"

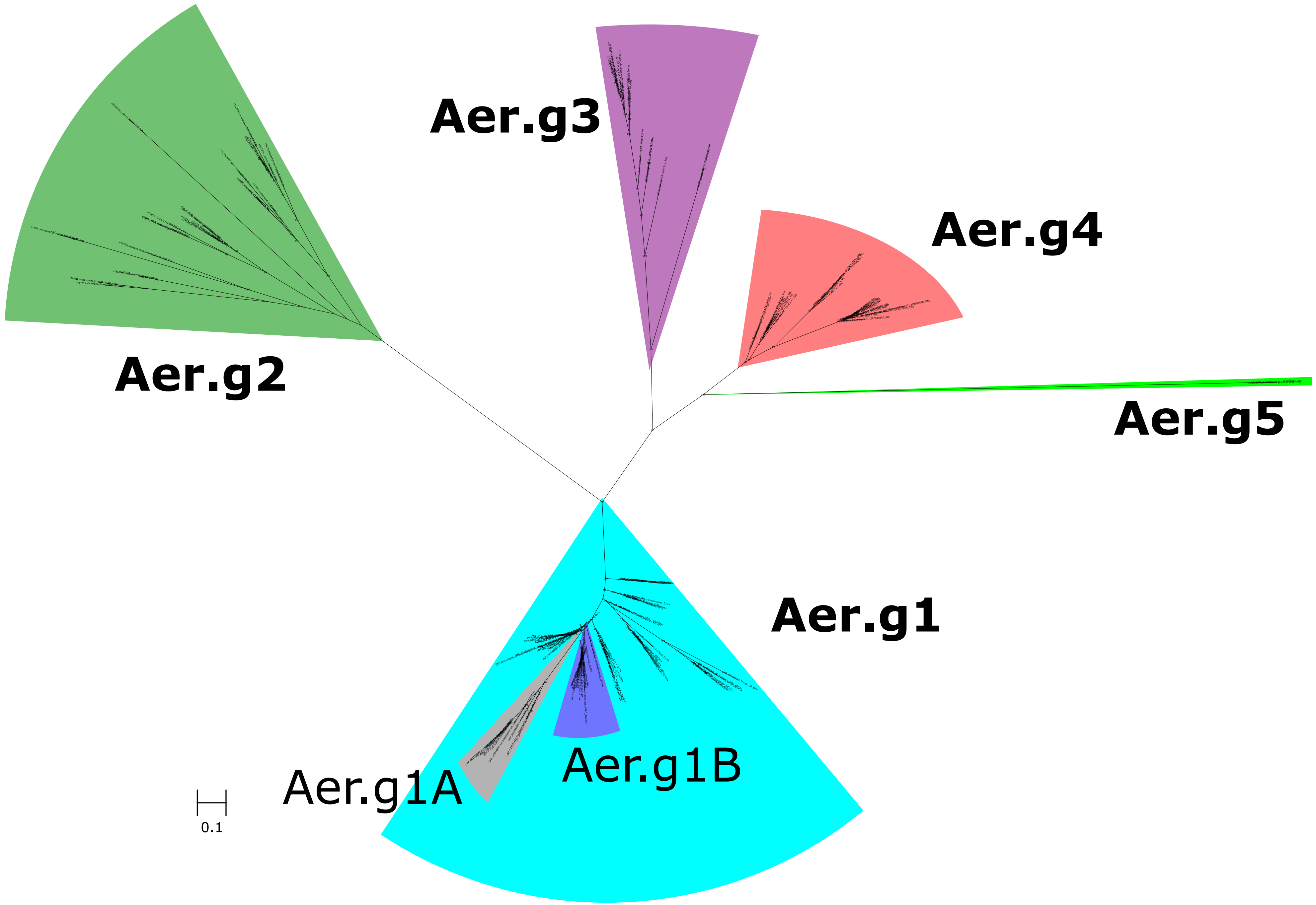


Supplementary Figure 1: Maximum-Likelihood unrooted consensus tree showing phylogenetic relationship between Aer protein sequences from *Pseudomonas* species. Sequences were grouped according to branching pattern and inspection of the alignment, and were confirmed in subsequent analysis, see following figures and text for details. Numbers along branches indicate bootstrap support values from 100 replicates. Scale bar indicates branch length equivalent to 0.1 AA substitutions per site.

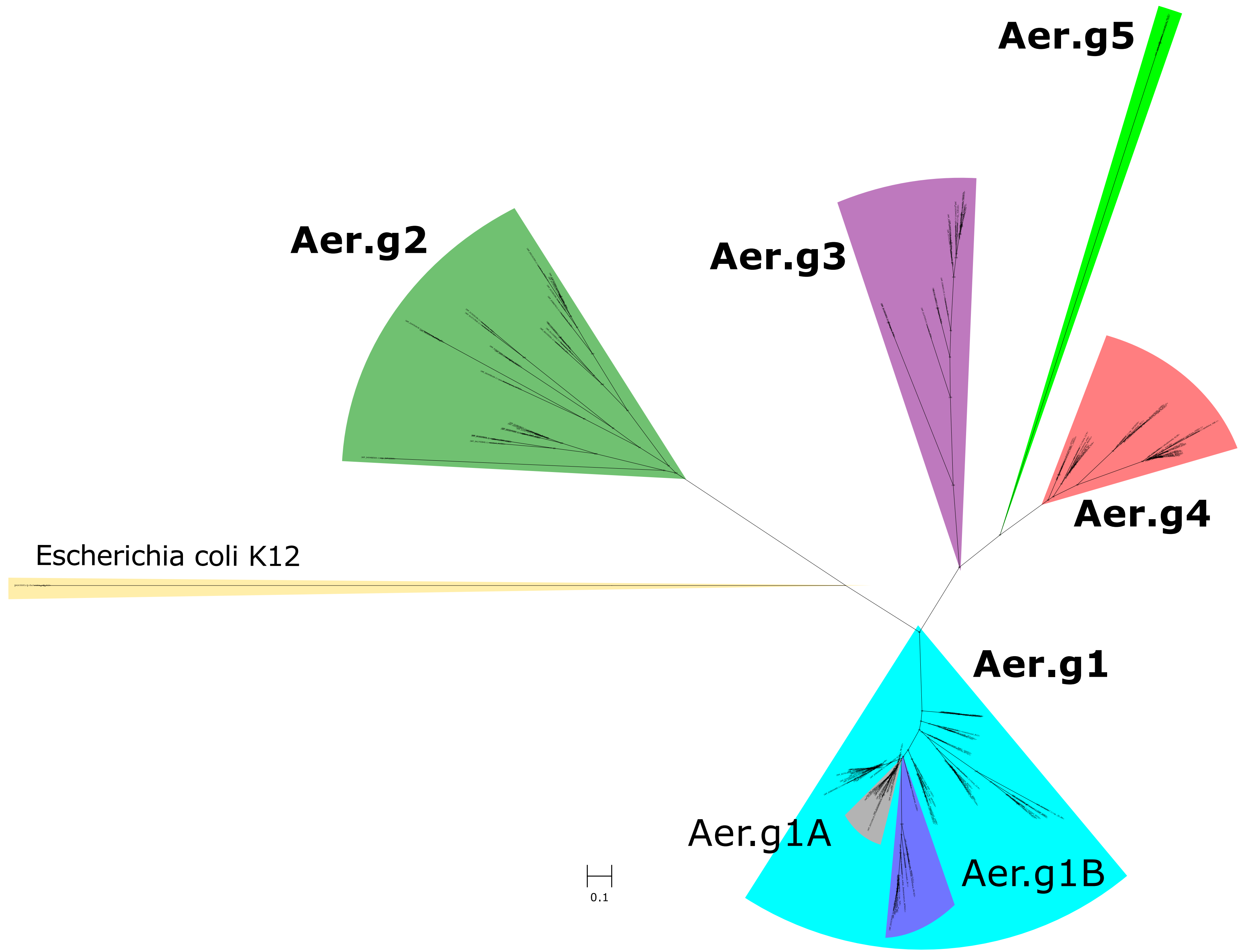


Supplementary Figure 2: Maximum-Likelihood unrooted consensus tree showing phylogenetic relationship between Aer protein sequences from *Pseudomonas* species, including Aer sequence from *E. coli*. Sequences were grouped according to branching pattern and inspection of the alignment, and were confirmed in subsequent analysis, see following figures and text for details. Numbers along branches indicate bootstrap support values from 100 replicates. Scale bar indicates branch length equivalent to 0.1 AA substitutions per site.

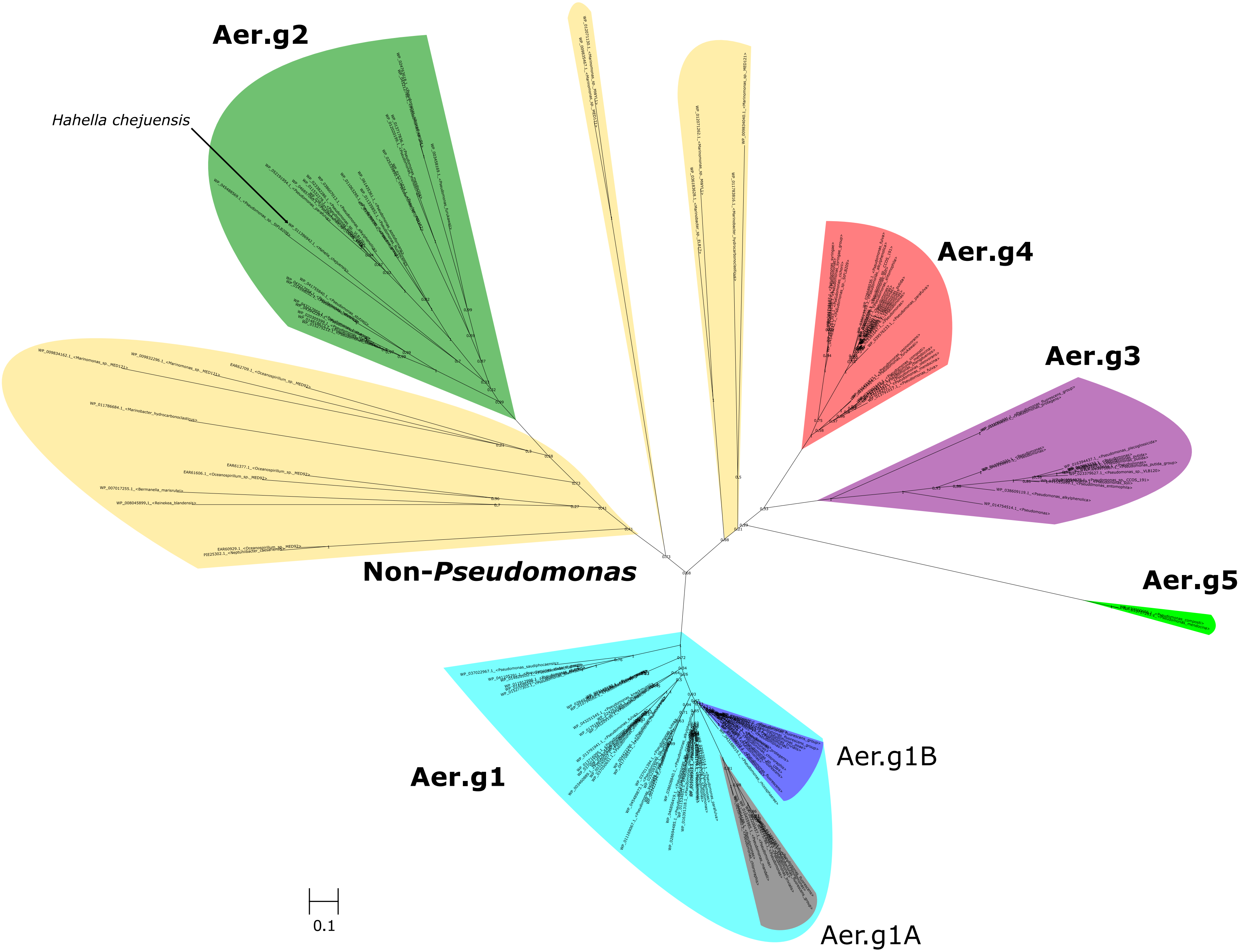


Supplementary Figure 3: Maximum-Likelihood consensus tree showing phylogenetic relationship between Aer protein sequences from *Pseudomonas* species, including probable Aer sequences from closely related non-*Pseudomonas* species. Sequences were grouped according to branching pattern and inspection of the alignment, and were confirmed in subsequent analysis, see following figures and text for details. Numbers along branches indicate bootstrap support values from 100 replicates. Scale bar indicates branch length equivalent to 0.1 AA substitutions per site.

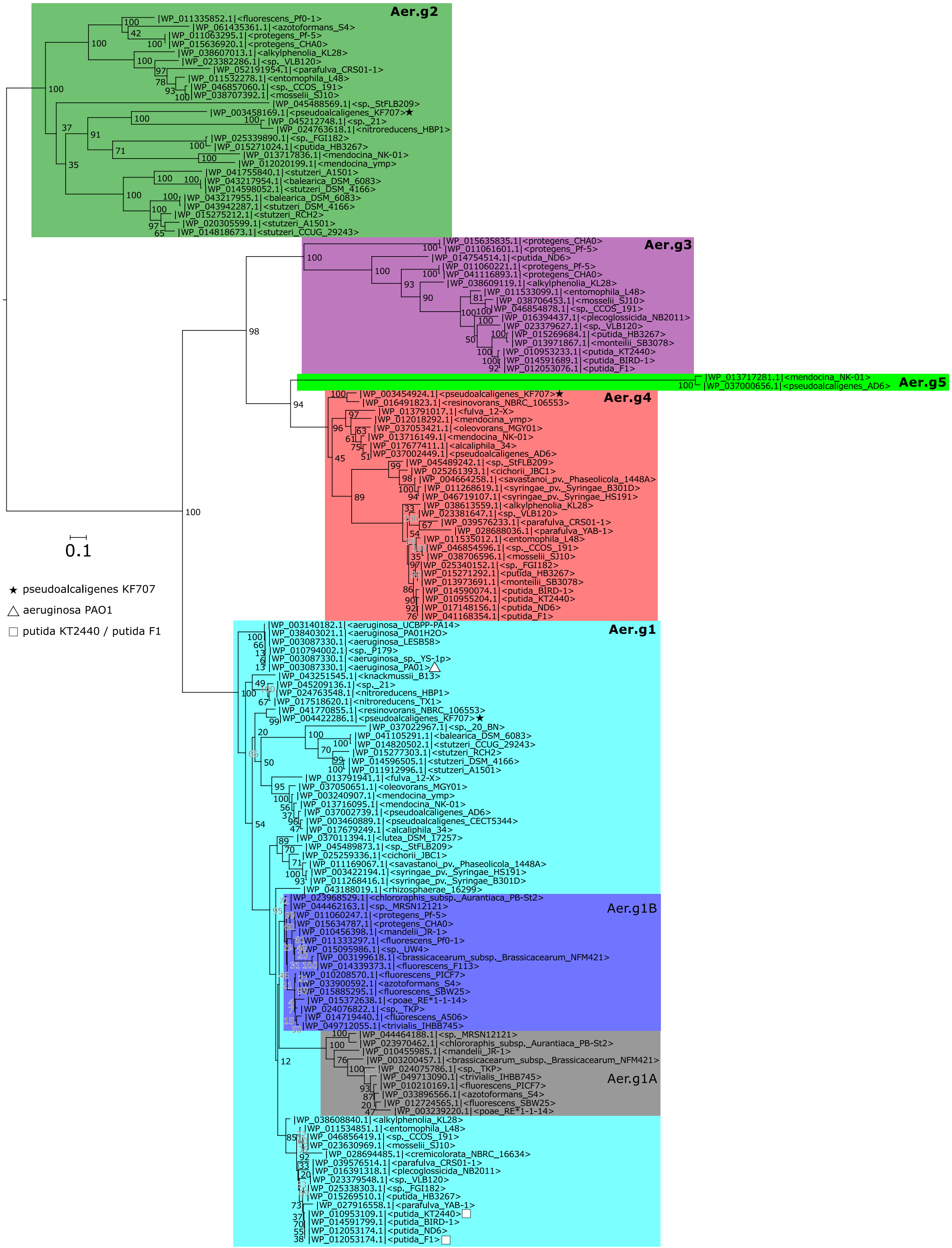


Supplementary Figure 3: Maximum-Likelihood consensus tree showing phylogenetic relationship between Aer protein sequences from *Pseudomonas* species. Sequences were grouped according to branching pattern and inspection of the alignment, and were confirmed in subsequent analysis, see following figures and text for details. The same characterized Aer sequences (triangle for *P. aeruginosa* PAO1, squares for *P. putida* F1 and KT2440), and sequences from P. pseudoalcaligenes KF707 (black stars) as Figure 1 are marked here. This tree is equivalent to the cladogram in Figure 1, but branches are not collapsed, and lengths and labels are maintained. The tree was generated unrooted, then the root placed based on comparison with unrooted trees containing either *E. coli* Aer or probable Aer homologs from closely related non-*Pseudomonas* species. See Supplementary Figures 1, 2 and 3 for details. Numbers along branches indicate bootstrap support values from 100 replicates. Scale bar indicates branch length equivalent to 0.1 AA substitutions per site.

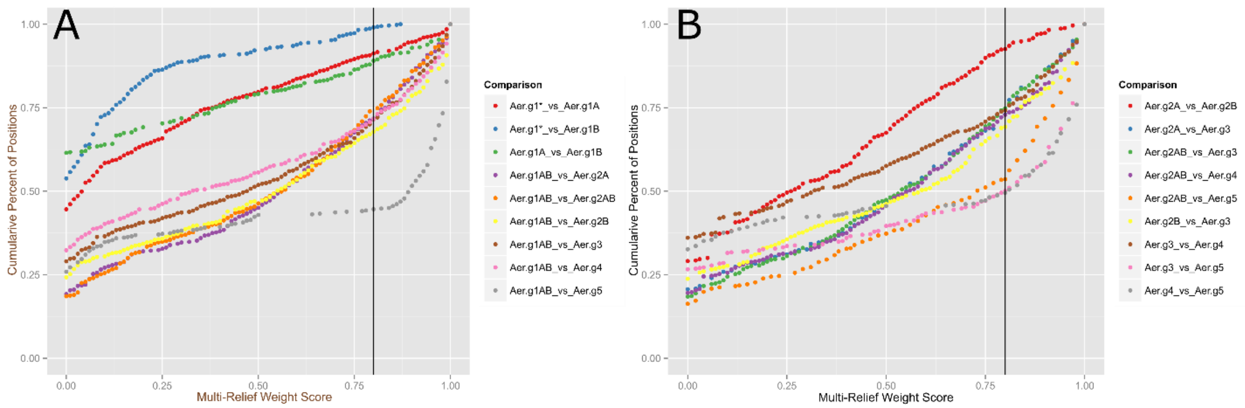


Supplementary Figure 5: Empirical cumulative distribution functions showing percent of amino acid positions at each multi-relief score value for comparisons of Aer homolog groups. Multi-relief weight scores ≥ 0.8 (black vertical line) indicate positions that are conserved within groups but not between groups. The point on the y-axis where each group comparison intersects with this line indicates the percentage of positions below the cutoff. Most group definitions result in ~>25% unique positions. Comparisons with group 5 show >50% unique positions (Aer.g5 only has two sequences). Intragroup comparisons between 1A/1B and 2A/2B show that these comparisons only result in <15% unique positions meaning their inclusion in their mother group is justified. Aer.g1* indicates Aer.g1 sequences not including Aer.g1A and Aer.g1B. Aer.g1AB includes all Aer.g1 sequences including Aer.g1A and Aer.g1B.

Sequence Harmony/Multi-Relief is a pair of algorithms that takes as input a pair of alignments and determines, for each AA position, whether that AA is conserved within each group, and whether it is divergent between the two groups, giving each AA position a score. A score of 1 indicates perfect within-group conservation and between-group divergence. Each pair of groups was compared in this fashion, then their results compared by determining what percent of AAs were conserved within a group but divergent between the groups. AA positions with a multi-relief weight score >0.8 were accepted as fitting these criteria (cutoff is based on the recommendations of the SHMR authors (25).

For each pair of groups a score was calculated for each amino acid position that indicated how conserved it was within groups and how divergent it was between groups. A score of 1 indicates perfect conservation within and perfect divergence between groups. Empirical cumulative distribution functions (ECDF) were plotted for each comparison and the percent of AAs above the 0.8 cutoff recommended by the SHMR authors was determined. As most comparisons resulted in 25% of AAs reaching this threshold, the two pairs of groups that, when compared, only had 10% of AAs above the cutoff, were deemed incorrect group assignments. These groups were merged and the process repeated to produce 5 groups that all had ~25% of AAs above the cutoff threshold. Group 5 was consistently excluded as it only contains 2 highly divergent sequences.

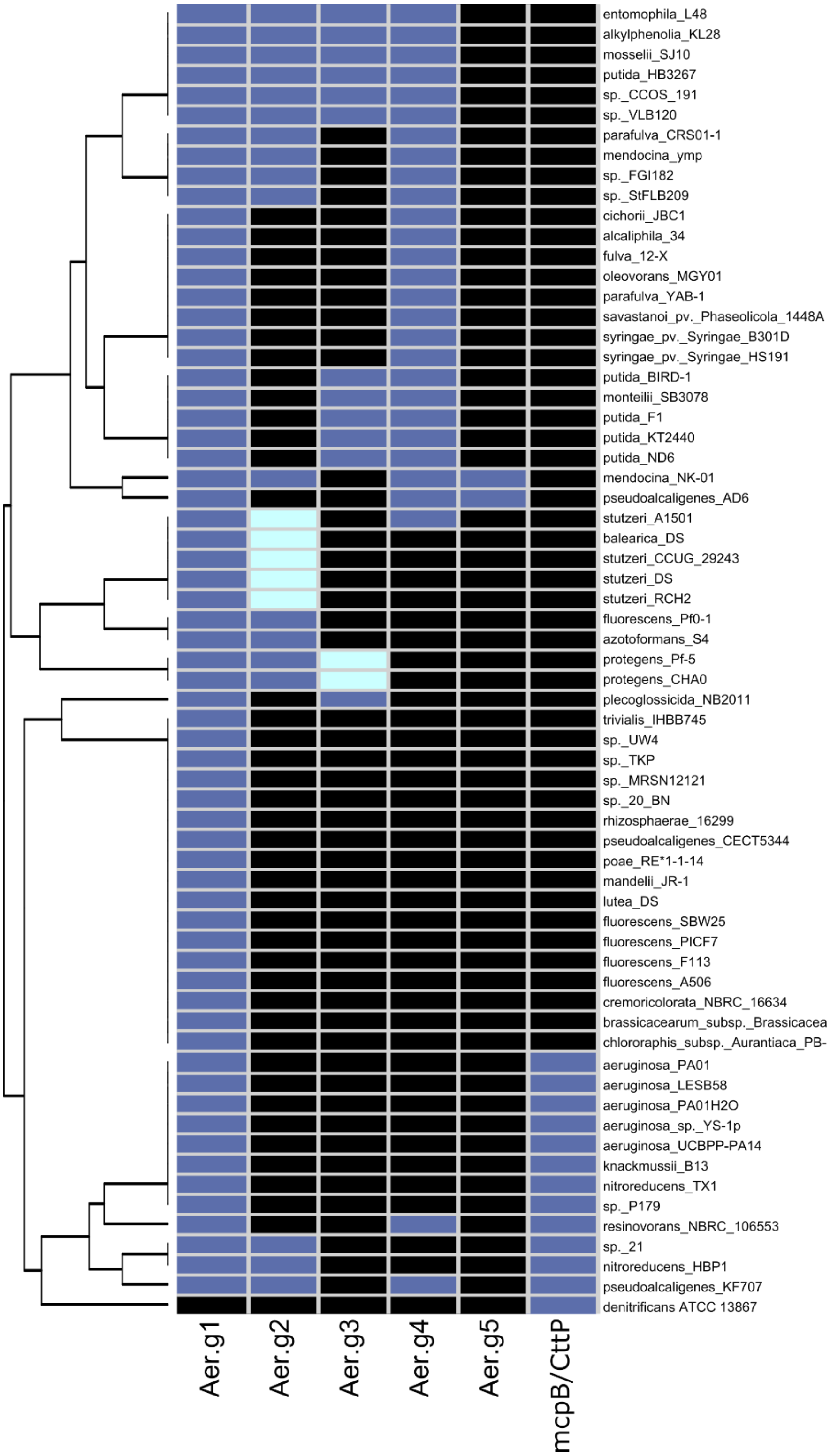


Supplementary Figure 6: Hierarchically clustered heatmap showing the presence and number of Aer homologs, McpB and Cttp in select *Pseudomonas* species. Species were clustered using the Bray-Curtis distance metric and average linkages. The number of homologs that each strain possess is indicated by the box colour: zero (black), one (blue), two (cyan).

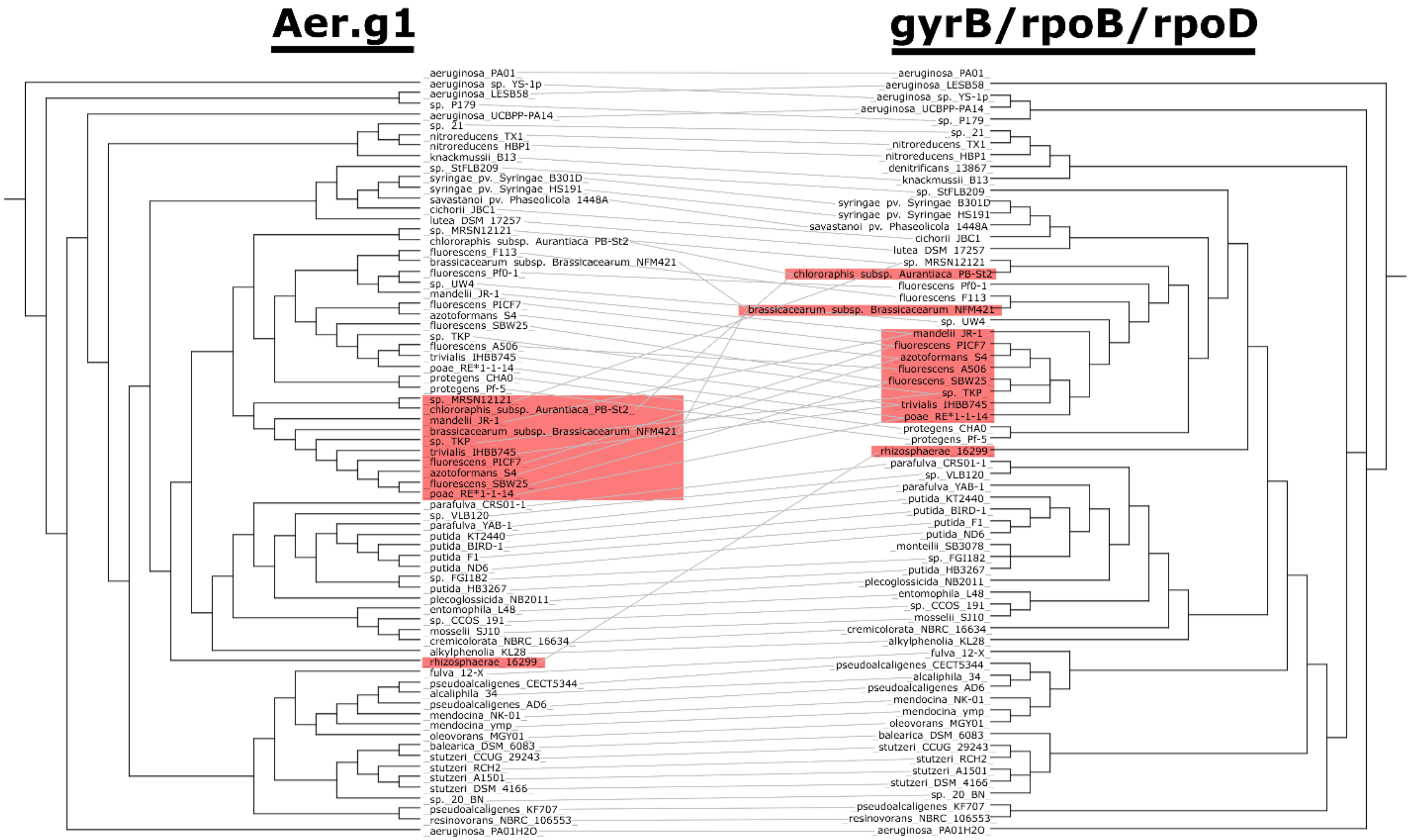


Supplementary Figure 7: Tanglegram showing tree matching between Aer.g1 subgroup tree and *Pseudomonas* genus phylogeny. Only species with a matching Aer sequence are included in the genus phylogeny. Matches highlighted in red indicate incongruity between the *gyrB/rpoB/rpoD* nucleotide sequence based tree and the Aer protein sequence based tree.

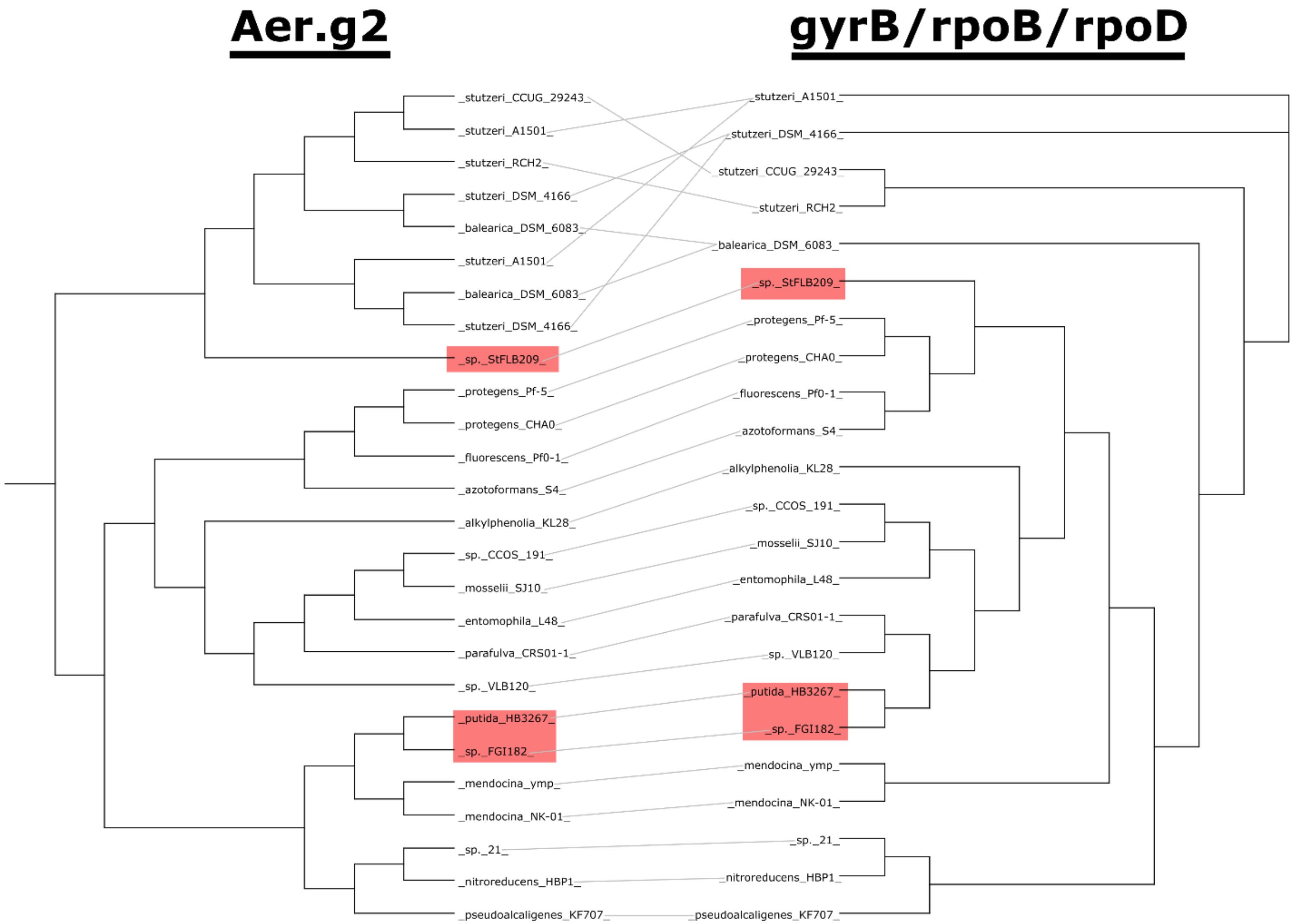


Supplementary Figure 8: Tanglegram showing tree matching between Aer.g2 subgroup tree and *Pseudomonas* genus phylogeny. Only species with a matching Aer sequence are included in the genus phylogeny. Matches highlighted in red indicate incongruity between the *gyrB/rpoB/rpoD* nucleotide sequence based tree and the Aer protein sequence based tree.

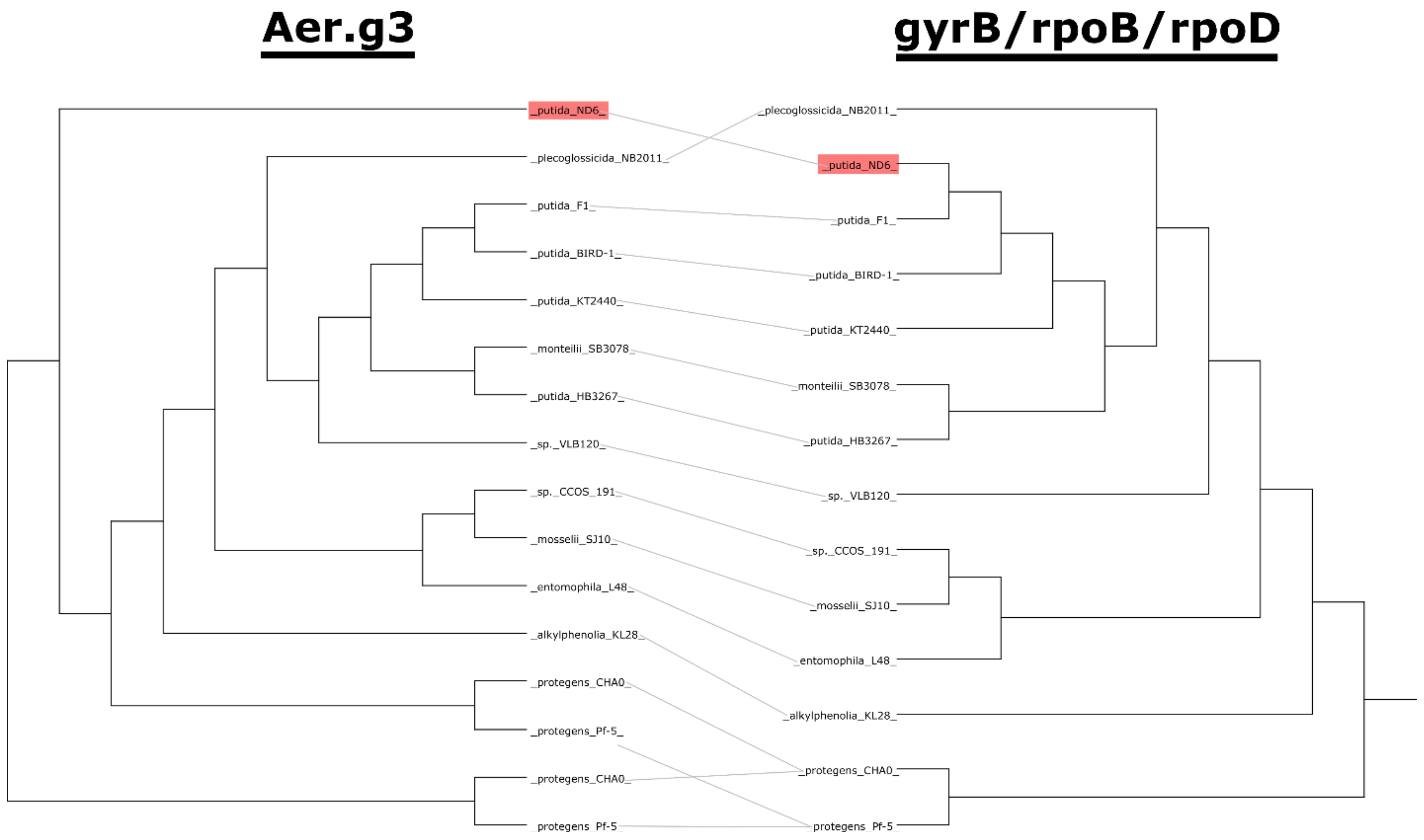


Supplementary Figure 9: Tanglegram showing tree matching between Aer.g3 subgroup tree and *Pseudomonas* genus phylogeny. Only species with a matching Aer sequence are included in the genus phylogeny. Matches highlighted in red indicate incongruity between the *gyrB/rpoB/rpoD* nucleotide sequence based tree and the Aer protein sequence based tree.

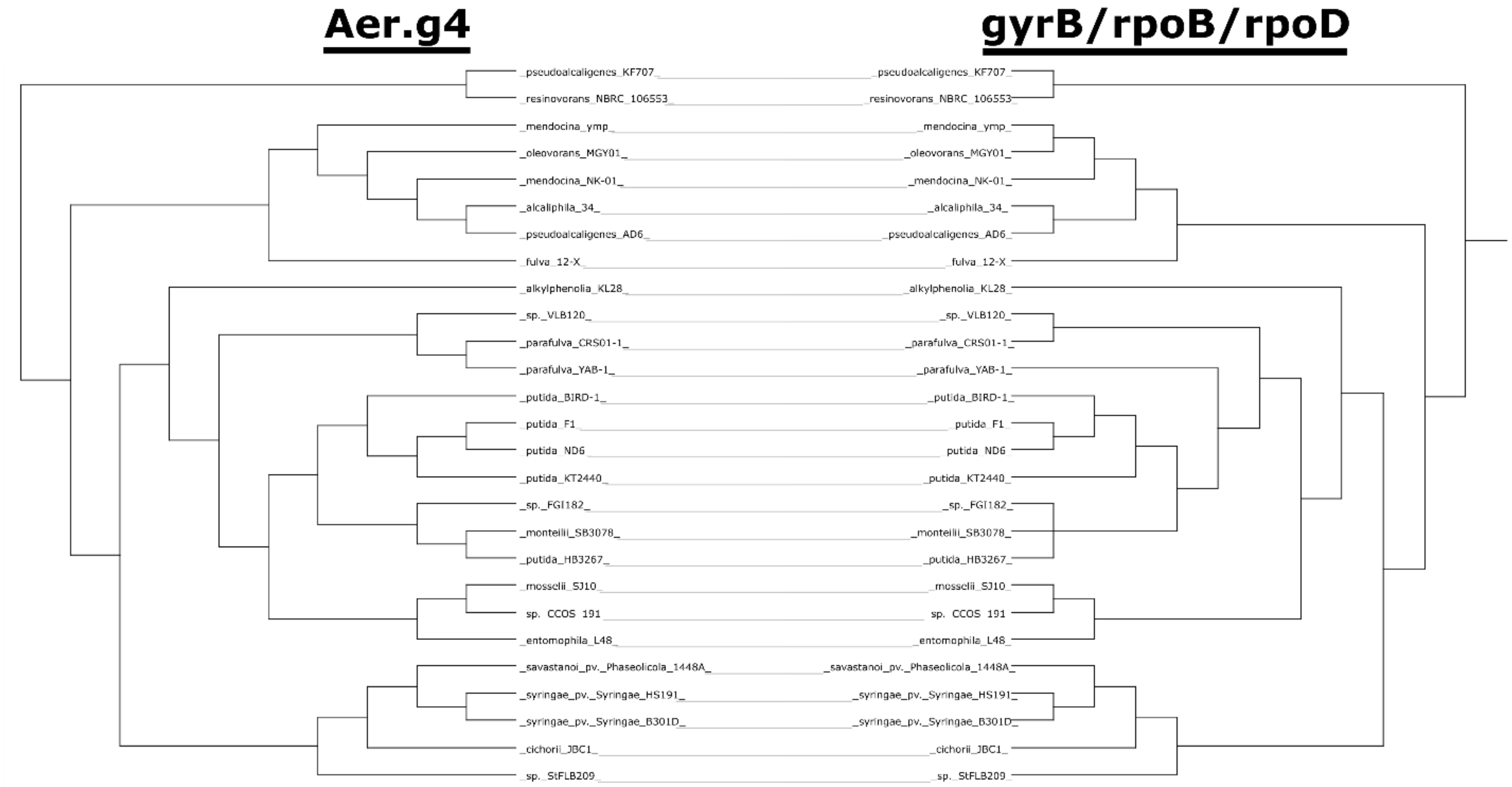


Supplementary Figure 10: Tanglegram showing tree matching between Aer.g1 subgroup tree and *Pseudomonas* genus phylogeny. Only species with a matching Aer sequence are included in the genus phylogeny.

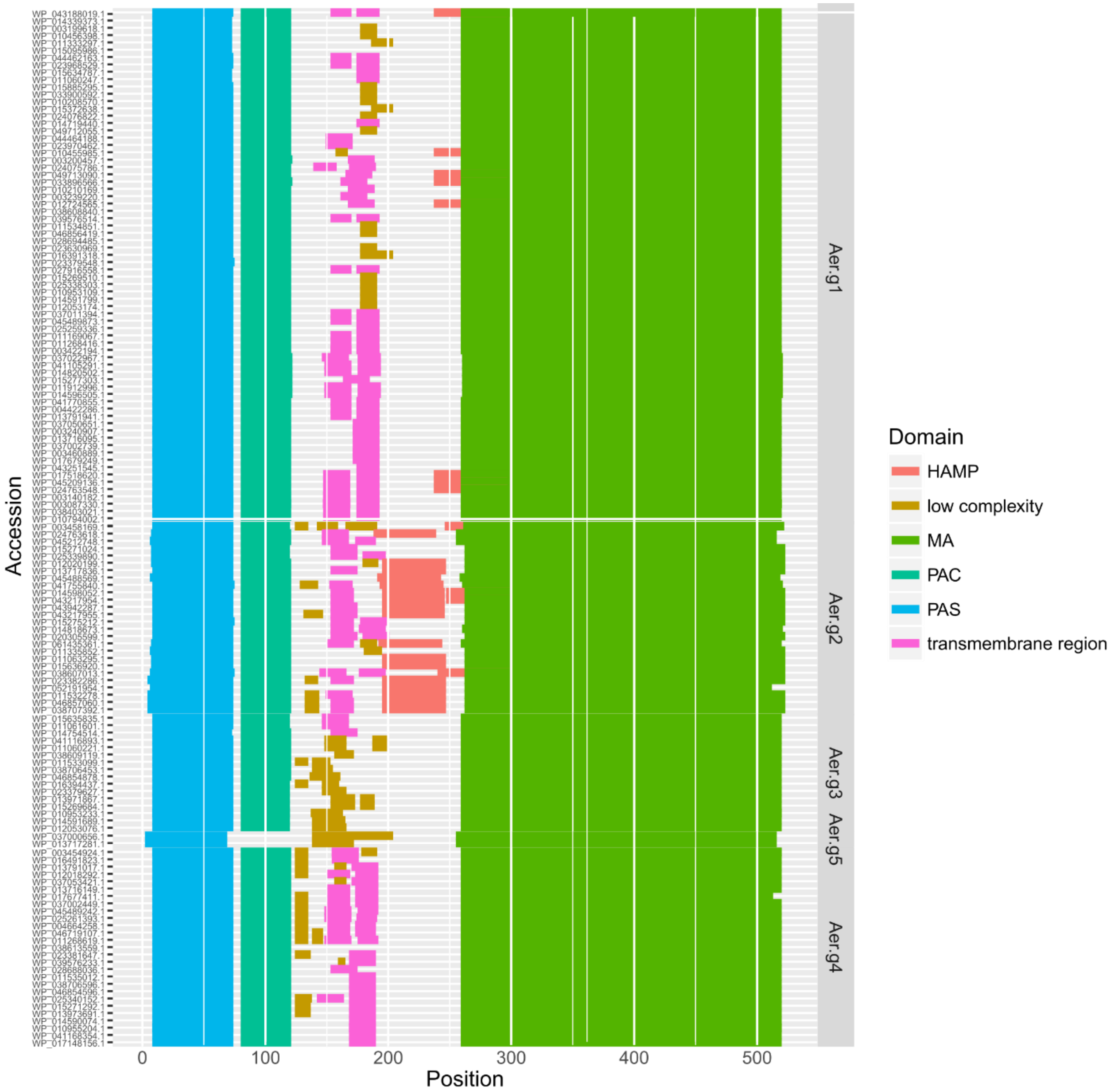


Supplementary Figure 11: Domain architecture of Aer sequences from select *Pseudomonas* species. Sequences were submitted to the SMART database and the results collected [1]. Domain start and end positions were used to mark coloured bars: PAS (Pern/Art/Sin, blue); PAC (PAS associated domain, teal); transmembrane region (pink); HAMP (Histidine kinase/adenylyl cyclases/methyl-accepting chemotaxis proteins/phosphatases, red); MA (methyl-accepting, green); low-complexity (brown). The methyl-accepting domain is also called the kinase control module or CheW/CheA interface.

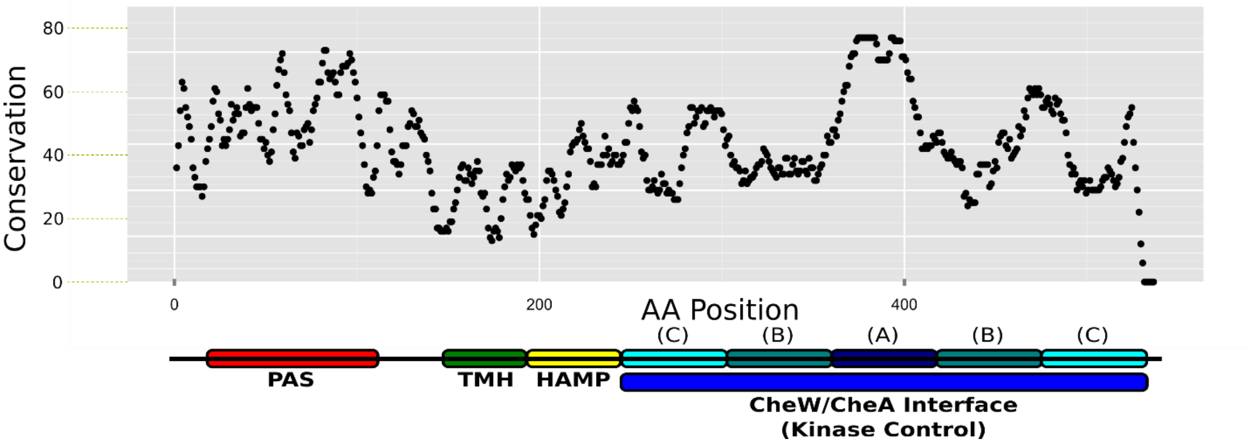


Supplementary Figure 12: Aer domain architecture and conservation across the entire length of all Aer sequences. Values were smoothed by taking the average of a position and the 3 preceding and following positions. Domains are PAS (Pern/Art/Sin) ligand binding (red), TMH (transmembrane helices, green), HAMP/Unique (in Aer.g2 HAMP, unique domains in other groups, yellow), CheW/CheA Interface also called Kinase Control (subdivided into A, signaling, dark blue; B, flexible bundle, teal; C, methylation, cyan).

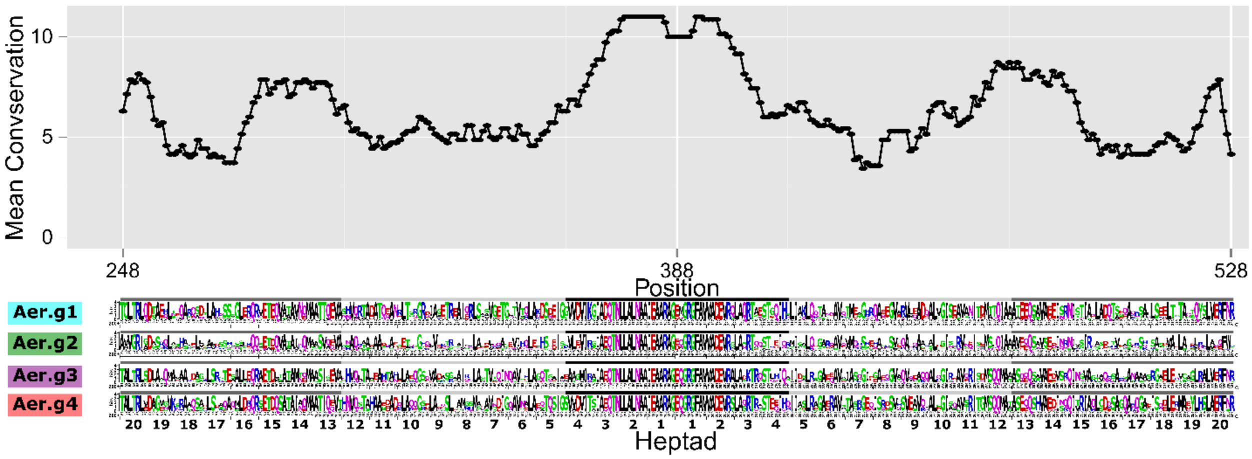


Supplementary Figure 13: Amino acid conservation of the cytoplasmic domain of Aer for all intergroup comparisons, and weblogos for each group. Conservation values were smoothed by taking the average of a position and the 3 preceding and following positions. The cytoplasmic domain is divided into 20 heptads, starting from the central glutamate counting outwards in each direction. Shading above the weblogos indicates the region (heptads 1-4, signaling; heptads 5-12, flexible bundle; heptads 13-20, methylation).

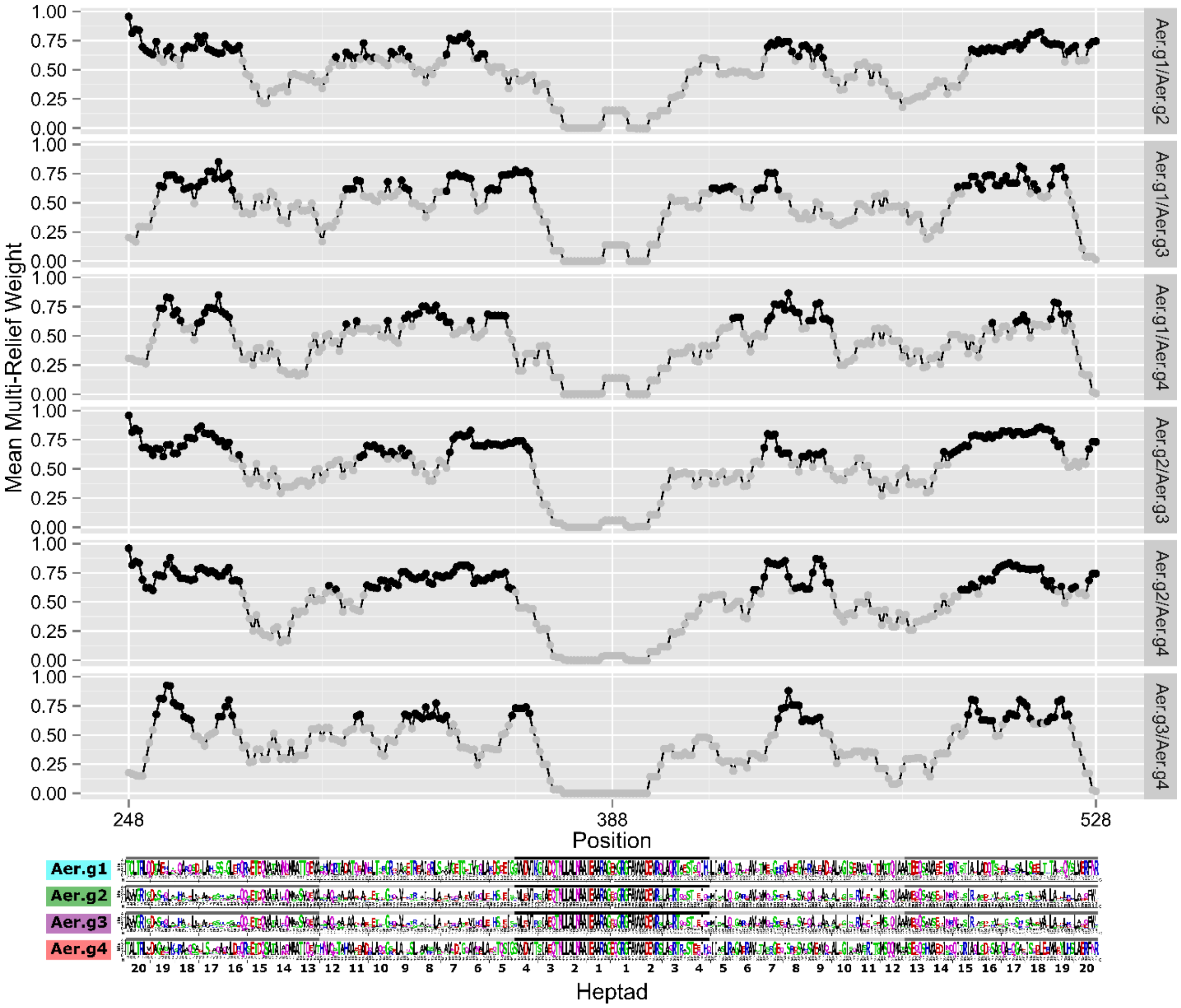


Supplementary Figure 14: Multi-relief scores of the cytoplasmic domain of Aer for all intergroup comparisons, and weblogos for each group. Multi-relief weight values were smoothed by taking the average of a position and the 3 preceding and following positions. Black points are above 0.6, grey below. The cutoff of 0.8 was relaxed due to the smoothing. Black regions indicate group unique regions. The cytoplasmic domain is divided into 20 heptads, starting from the central glutamate counting outwards in each direction. Shading above the weblogos indicates the region (heptads 1-4, signaling; heptads 5-12, flexible bundle; heptads 13-20, methylation).

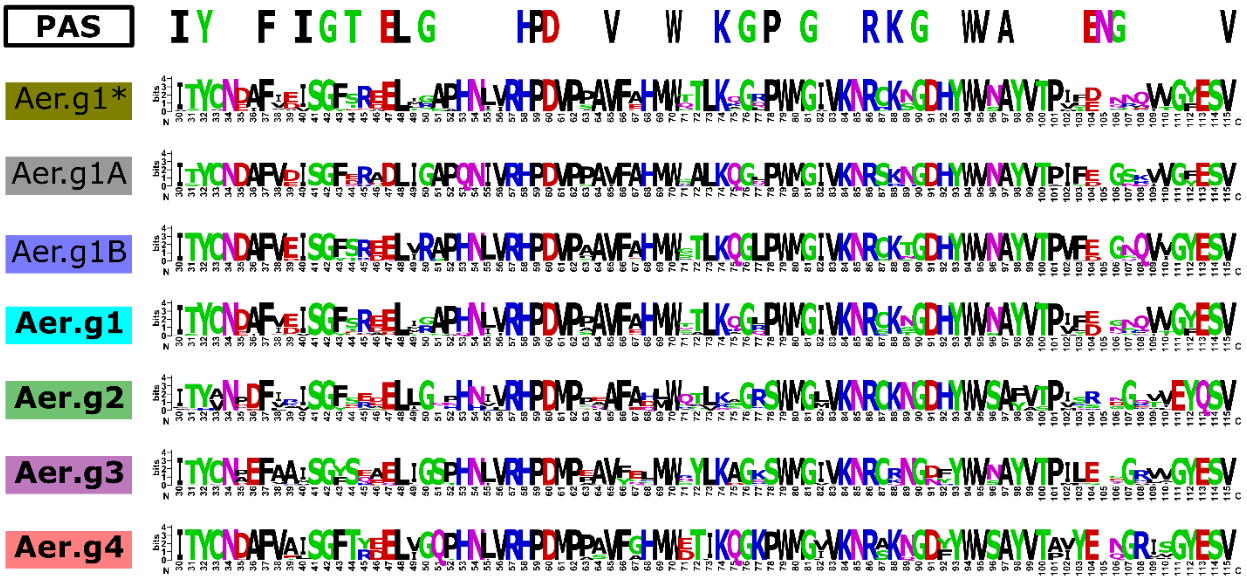


Supplementary Figure 15: Weblogos of each Aer group's PAS domain. Aer.g1* indicates Aer.g1 sequences not including Aer.g1A and Aer.g1B. Aer.g1 includes all Aer.g1 sequences including Aer.g1A and Aer.g1B. Characteristic PAS domain residues were obtained from the SMART database.

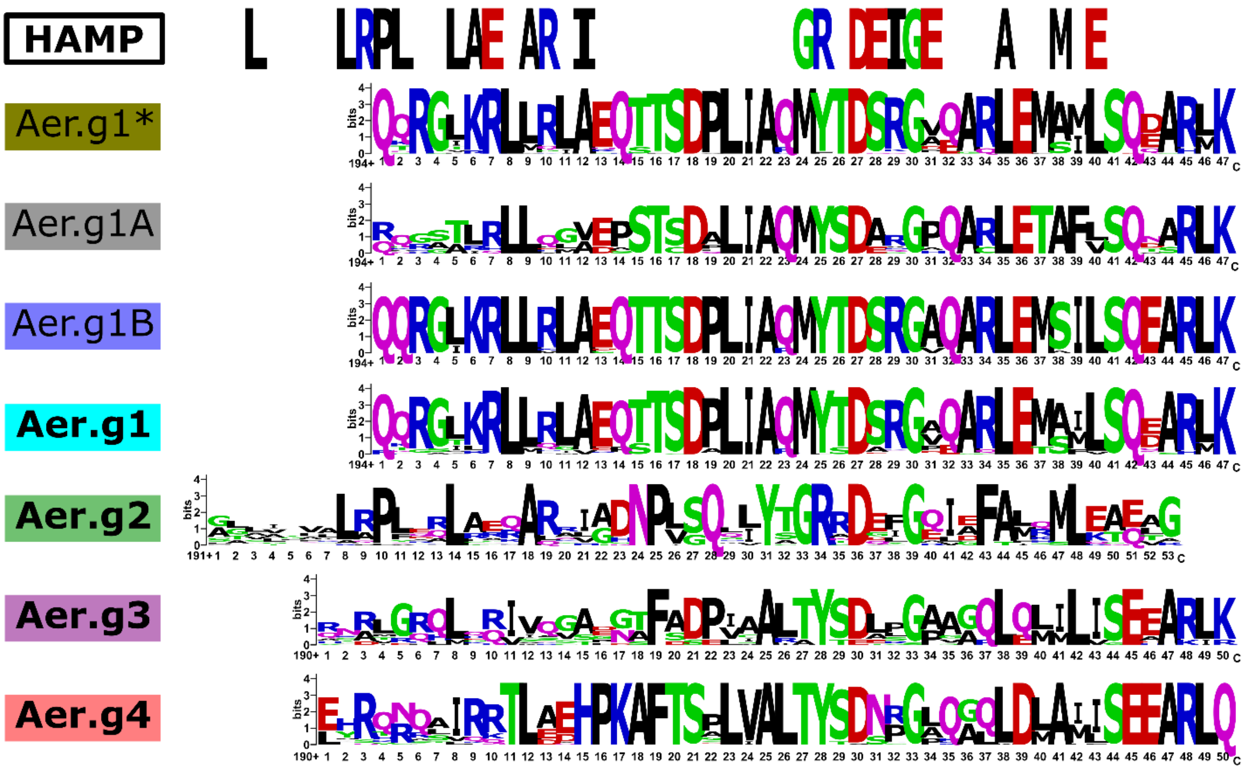


Supplementary Figure 16: Weblogos of Aer group specific region in between end of transmembrane domain and start of cytoplasmic domain heptads. Start of cytoplasmic domain was based on the number of matching heptads on either side of the central glutamate residue (Aer.g1, 388; Aer.g2 394; Aer.g3, 394; Aer.g4 391). The end of the transmembrane domain was determined by submitting the consensus sequence of each group to THMMR. Weblogos were aligned based on the shared aspartate residue

(Aer.g1, 221; Aer.g2 227; Aer.g3, 220; Aer.g4 220). Characteristic HAMP domain residues were obtained from the SMART database. This figure is the same as Figure 5, only it includes the Aer.g1 subdivisions. Aer.g1* indicates Aer.g1 sequences not including Aer.g1A and Aer.g1B. Aer.g1 includes all Aer.g1 sequences including Aer.g1A and Aer.g1B. This figure is equivalent to Figure 5, but includes the subdivisions of Aer.g1.

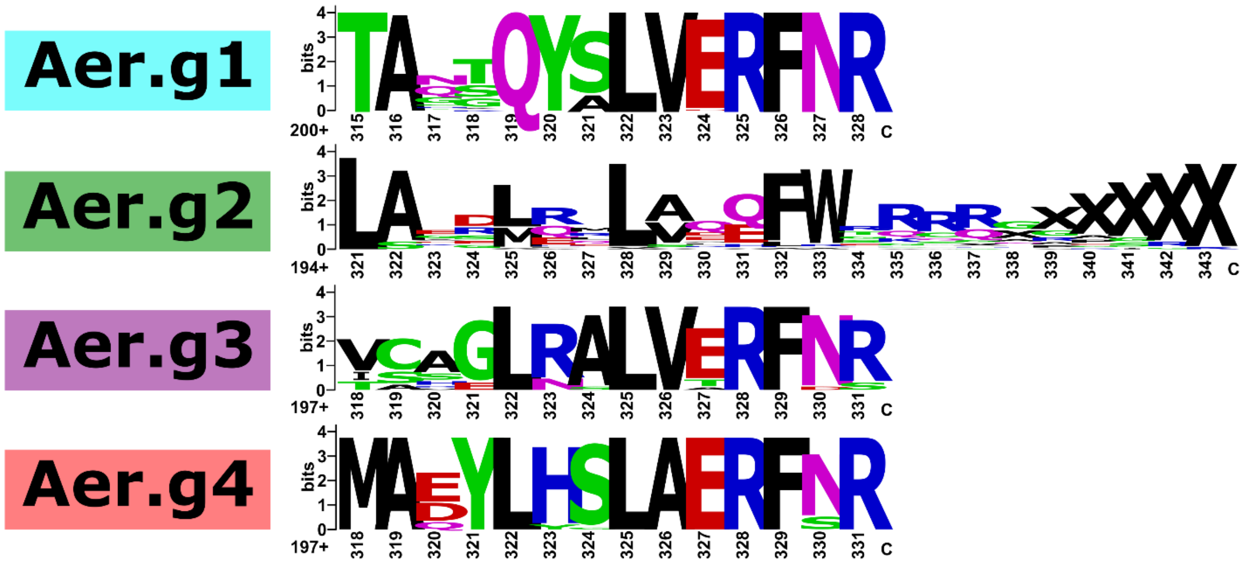


Supplementary Figure 17: Weblogos of Aer C-terminal region. Except for Aer.g2, this constitutes a zoomed in region shown in Supplementary Figure 12. X denotes NO amino acid in that position.

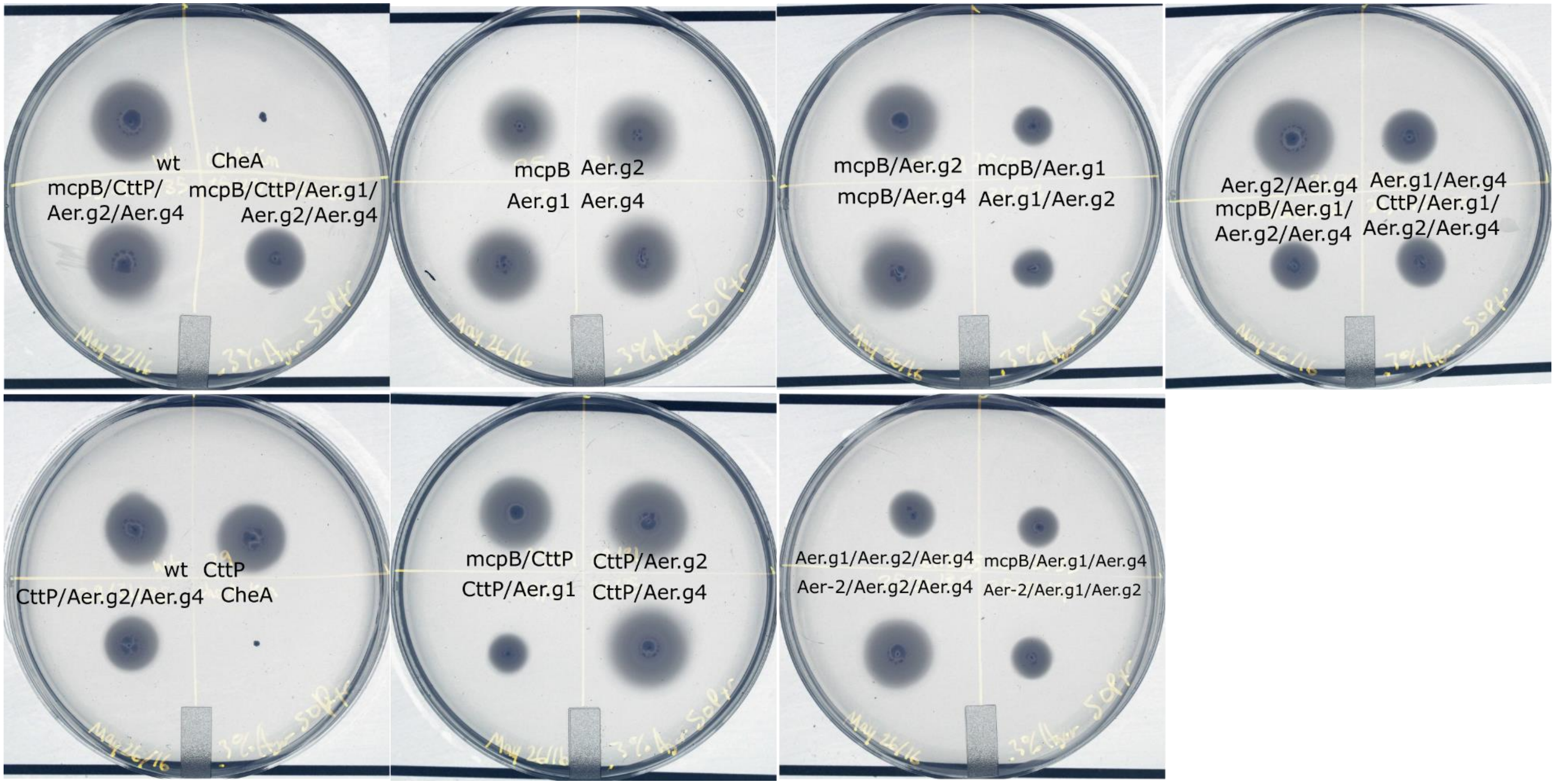


Supplementary Figure 18: Photographs of energy-taxis swim plates of *P. pseudoalcaligenes* strains with deletions in *mcpB*, *cttP* and *aer* homologs after 24h growth at 30°C in 50mM pyruvate. Colours have been inverted to emphasize contrast.

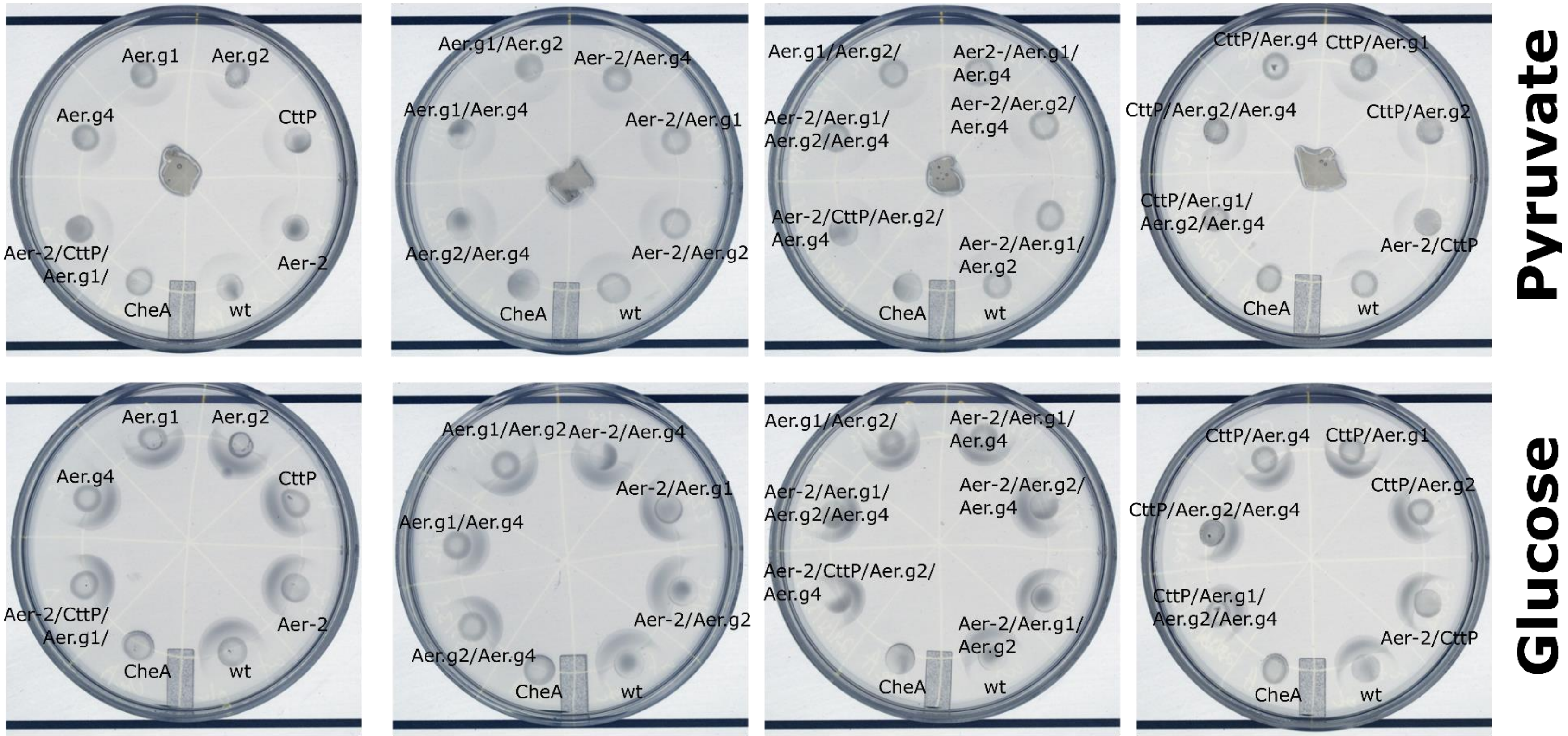


Supplementary Figure 19: Chemotaxis swim assays of *P. pseudoalcaligenes* strains with deletions in *mcpB, cttP* and *aer* homologs. Strains were grown overnight, concentrated then spotted on minimal salts plates containing 0.3% agar. Either 50mM pyruvate in 1.5% agar or crystals of glucose were placed in the centre of the plates. Photographs were taken after 24h. Colours have been inverted to emphasize contrast.

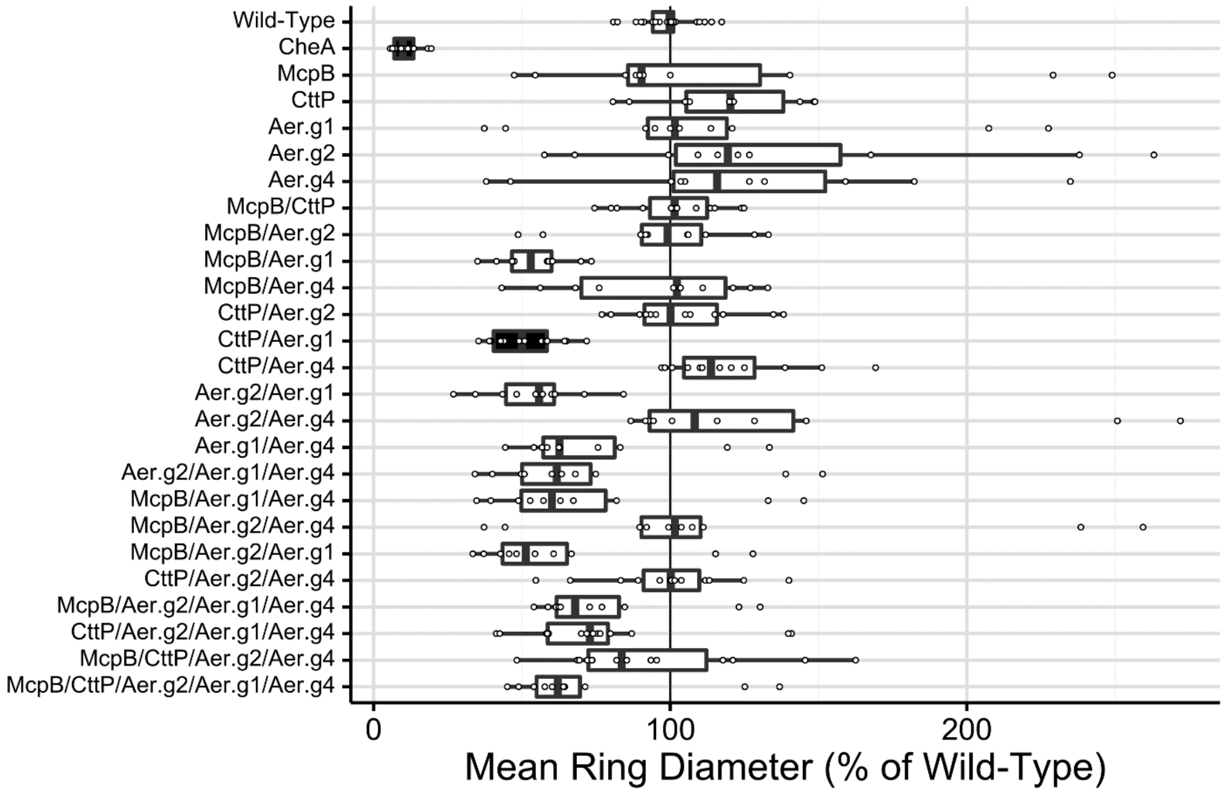


Supplementary Figure 20: Mean normalized energy-taxis diameters of strains of *P. pseudoalcaligenes* KF707 with deletions of *mcpB, cttP* and *aer* homologs in 50mM succinate. Boxes filled in dark-grey indicate significant differences from the wild-type based on Tukey's Honest Significant Differences test with a confidence value of 0.95. Values were normalized to the wild-type within each experiment at 24 and 48 h, allowing datapoints from both times to be combined. Wild-type strains were normalized to the mean of technical replicates within each experiment. The *cheA::KmR* mutant was not grown with antibiotic present.

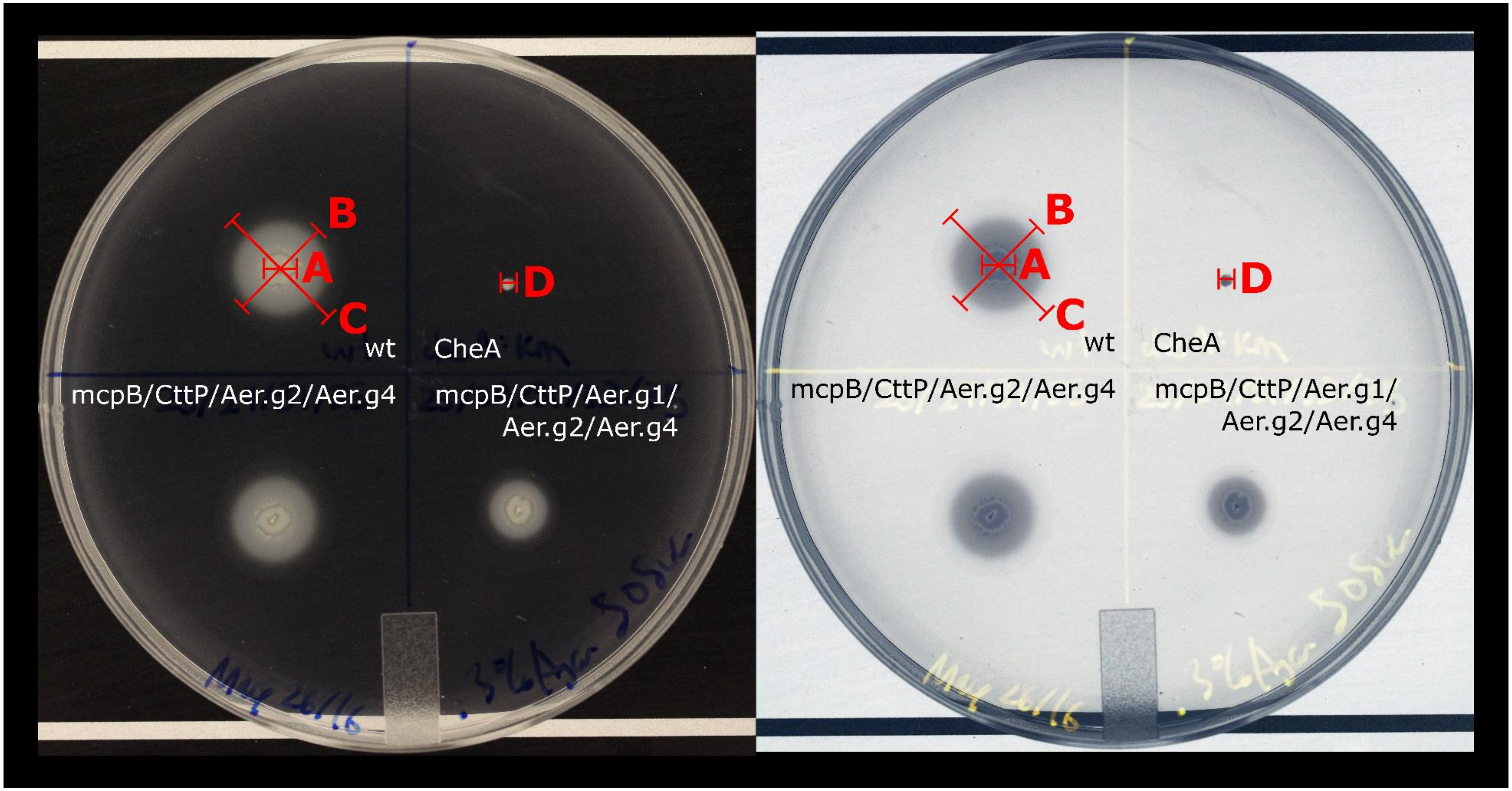


Supplementary Figure 21: Diagram showing how energy-taxis phenotypes were measured. Left is the actual image, right is the same image with colours inverted. This example plate shows the growth of 4 strains after 24h in minimal salts medium, 0.3% agar, 50mM succinate. Strains were inoculated into the plate on a sterile needle. Three different migration horizons are visible: the edge of dense growth (A), the intermediate colony edge (B), the edge of swimming cells (C). In the CheA mutant, there is no separation of horizons as the strain is non chemotactic (D). The diameter as defined by C that crossed through the inoculation center was used to measure the energy taxis diameter discussed in further figures.

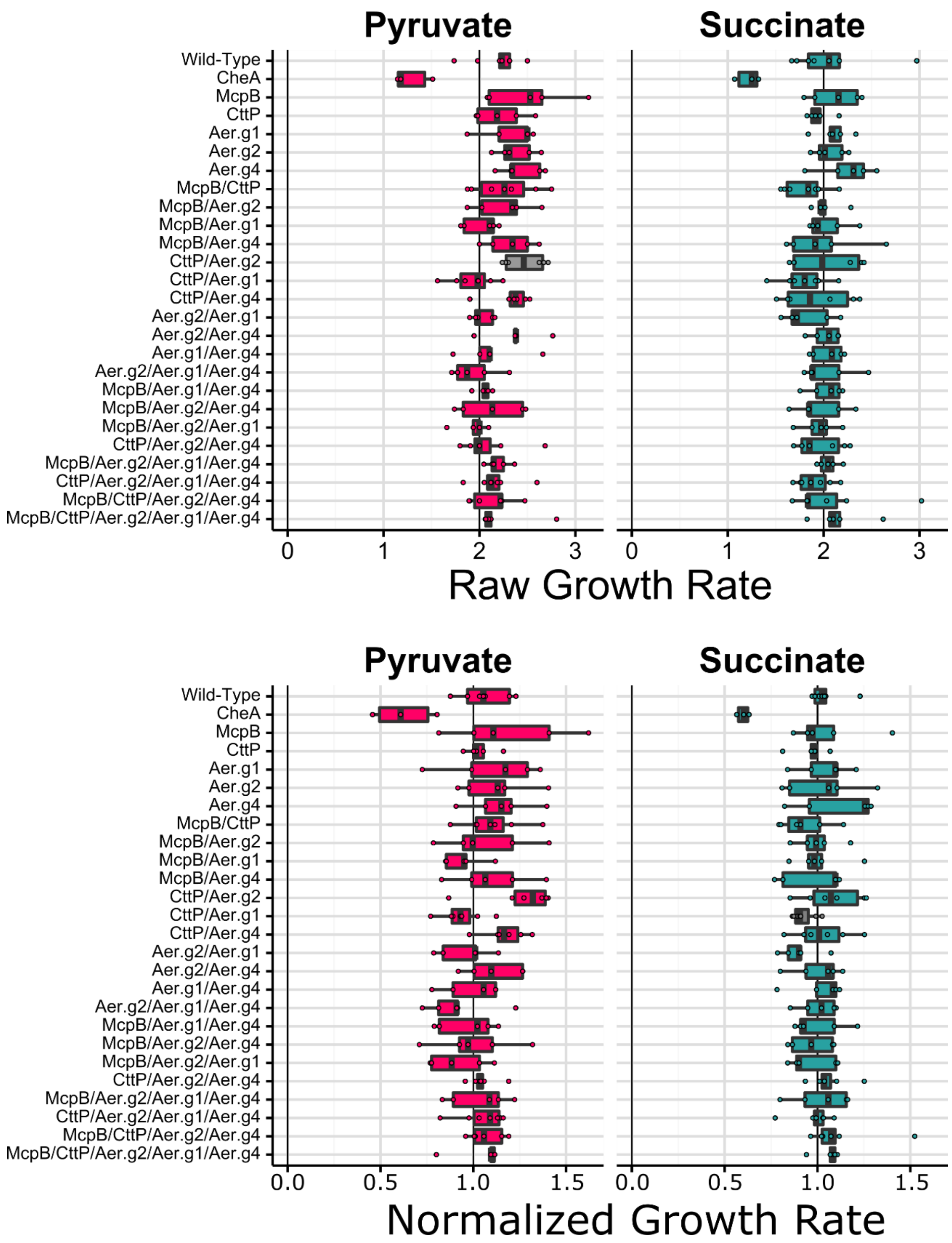


Supplementary Figure 22: Mean raw and normalized growth rates of energy-taxis swim-diameters of strains of *P. pseudoalcaligenes* KF707 with deletions of *mcpB, cttP* and *aer* homologs. Growth rates were calculated by dividing the diameter at 48h by the diameter at 24h. Normalized rates were calculated from diameters that were normalized to the wild-type for each experiment at each time point. Stars indicate significant differences from the corresponding wild-type according to Tukey's Honest Significant Differences test. Values represent results from at least 3 experimental replicates.

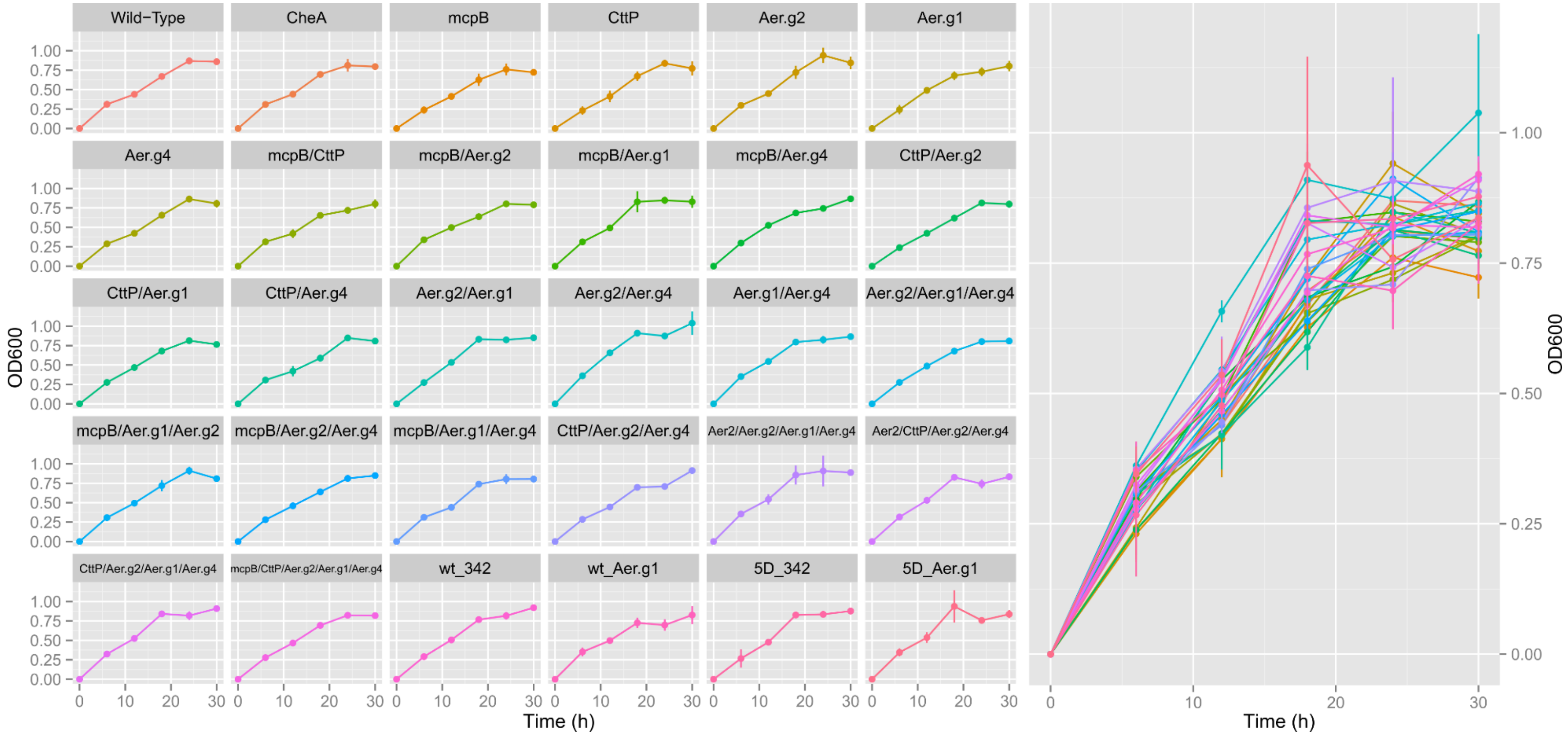


Supplementary Figure 23: Growth *P. pseudoalcaligenes* KF707 strains with deletions of Aer homologs, CttP and McpB. Growth was performed in a microtiter plate in minimal salts medium with 10mM pyruvate as the growth substrate. Growth of the wild-type and quintuple mutant (McpB/CttP/Aer.g1/Aer.g2/Aer.g4) with the empty vector pSEVA342 and pSEVA342_Aer.g1 was also assayed. Values represent the average of 2 biological replicates from 2 experimental replicates (4 total replicates). Lines at points indicates standard error.

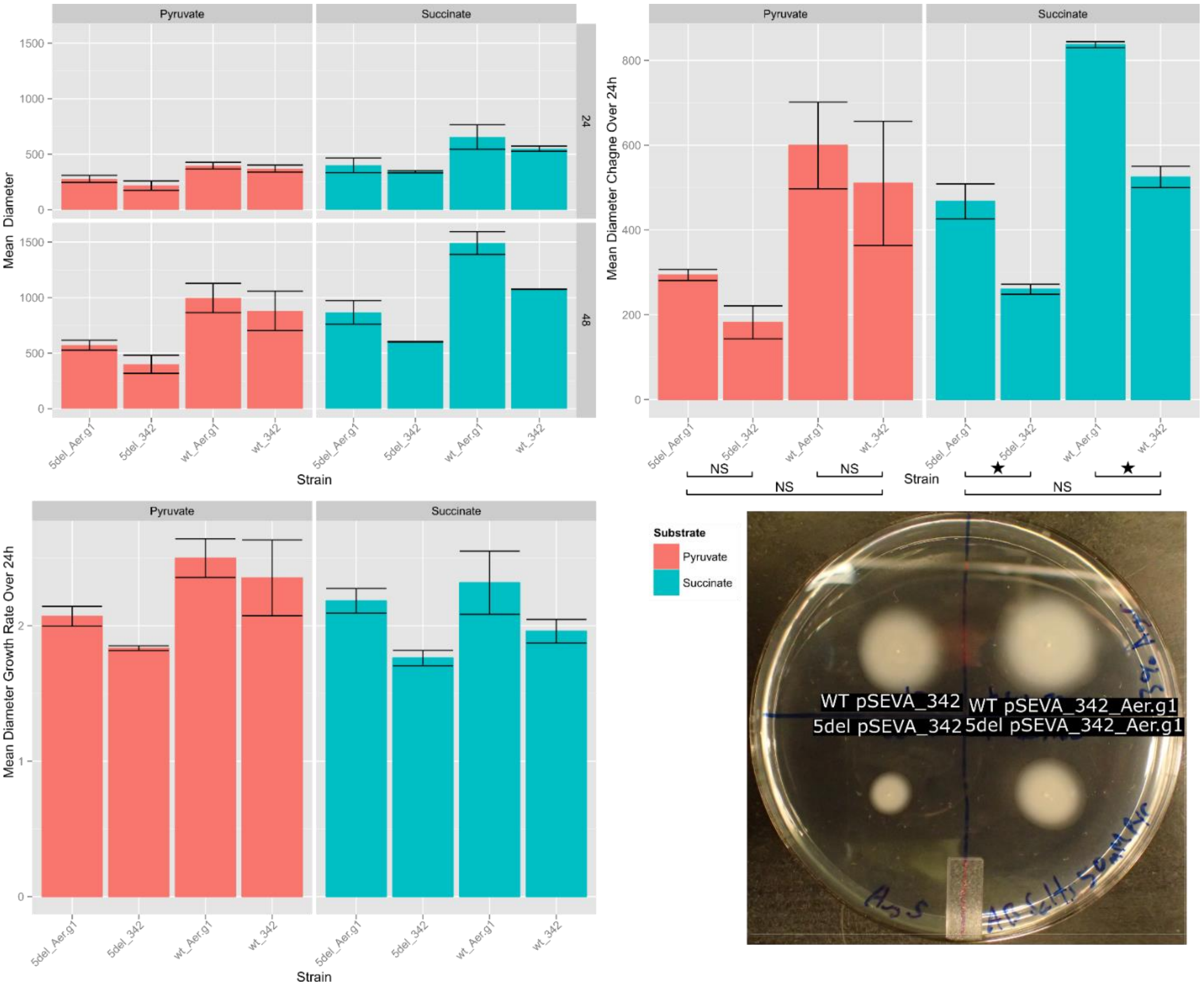


Supplementary Figure 24: Mean energy-taxis diameters, change of diameters over 24h and rate of growth of diameters of strains of *P. pseudoalcaligenes* KF707 wild-type and quintuple mutant strains carrying Aer.g1 complementation plasmids. Growth rates were calculated by dividing the diameter at 48h by the diameter at 24h. Change is the difference between 24h and 48h. Stars indicate significant differences according to Tukey's Honest Significant Differences test. Values represent results from 2 experimental replicates. Strains were carrying either pSEVA_342 (empty vector control) or p_SEVA_342 with Aer.g1. 5del indicates quintuple mutant with deletion of Aer2/CttP/Aer.g2/Aer.g1/Aer.g4. Photograph in bottom right was taken after 24h of growth.

Supplementary Table 1: Genes immediately adjacent to *aer* genes and presence of mobile elements. See supplementary data.

Supplementary Table 2: Frequency of association of mobile elements within 5kb of *aer* genes. Summarized from supplementary Table 1.

| Aer Group | Repeats | Transposase | Integrase |
|---|---|---|---|
| aer.g1 | 20/73 | 2/73 | 0/73 |
| aer.g2 | 4/26 | 1/26 | 0/26 |
| aer.g3 | 1/16 | 1/16 | 1/16 |
| aer.g4 | 4/27 | 2/27 | 1/27 |
| aer.g5 | 0/2 | 0/2 | 0/2 |
| Total | 29/144 | 6/144 | 2/144 |

Supplementary Table 3: Tukey Honest Significant Differences results comparing differences in normalized energy-taxis diameter in pyruvate or succinate plates. HSD test compared all pairs of strains, only comparisons to the wild-type are presented here. P values were computed using a 0.95 confidence value, those below 0.05 were taken as significant.

| Comparison | Pyruvate Difference | Pyruvate Adjusted p Value | Succinate Difference | Succinate Adjusted p Value |
|---|---|---|---|---|
| Wild-Type-Aer.g1 | 0.044557106 | 1 | -0.15022 | 1 |
| Wild-Type-Aer.g1/Aer.g4 | 0.316929667 | **0.001584542** | 0.240355 | 0.995568 |
| Wild-Type-Aer.g2 | -0.130260185 | 0.994474052 | -0.37755 | 0.470077 |
| Wild-Type-Aer.g2/Aer.g1 | 0.416623963 | **6.99E-07** | 0.449853 | 0.123956 |
| Wild-Type-Aer.g2/Aer.g1/Aer.g4 | 0.41531338 | **7.83E-07** | 0.259256 | 0.985363 |
| Wild-Type-Aer.g2/Aer.g4 | -0.107657177 | 0.999855723 | -0.38801 | 0.404151 |
| Wild-Type-Aer.g4 | -0.080272111 | 0.999999891 | -0.23695 | 0.996521 |
| Wild-Type-Aer2 | 0.06079056 | 1 | -0.18355 | 0.99998 |
| Wild-Type-Aer2/Aer.g1 | 0.423034811 | **3.99E-07** | 0.452405 | 0.116907 |
| Wild-Type-Aer2/Aer.g1/Aer.g4 | 0.348397359 | 0.000171831 | 0.267403 | 0.977193 |
| Wild-Type-Aer2/Aer.g2 | -0.028743339 | 1 | 0.025494 | 1 |
| Wild-Type-Aer2/Aer.g2/Aer.g1 | 0.424269068 | **3.58E-07** | 0.35864 | 0.594415 |
| Wild-Type-Aer2/Aer.g2/Aer.g1/Aer.g4 | 0.358356796 | **8.13E-05** | 0.203102 | 0.999816 |
| Wild-Type-Aer2/Aer.g2/Aer.g4 | 0.172011243 | 0.810645941 | -0.19209 | 0.999944 |
| Wild-Type-Aer2/Aer.g4 | 0.00831534 | 1 | 0.050637 | 1 |
| Wild-Type-Aer2/CttP | -0.05951297 | 0.999999999 | -0.02338 | 1 |
| Wild-Type-Aer2/CttP/Aer.g2/Aer.g1/Aer.g4 | 0.396996279 | 3.72E-06 | 0.326159 | 0.792546 |
| Wild-Type-Aer2/CttP/Aer.g2/Aer.g4 | 0.063072257 | 0.999999997 | 0.012354 | 1 |
| Wild-Type-CheA | 0.87948192 | 0 | 0.88287 | **3.16E-10** |
| Wild-Type-CttP | -0.139877304 | 0.983128982 | -0.19063 | 0.999953 |
| Wild-Type-CttP/Aer.g1 | 0.459331863 | **4.31E-11** | 0.480318 | **0.011177** |
| Wild-Type-CttP/Aer.g2 | -0.025195491 | 1 | -0.04643 | 1 |
| Wild-Type-CttP/Aer.g2/Aer.g1/Aer.g4 | 0.38554092 | **1.60E-07** | 0.22296 | 0.991638 |
| Wild-Type-CttP/Aer.g2/Aer.g4 | 0.015579387 | 1 | 0.001138 | 1 |
| Wild-Type-CttP/Aer.g4 | -0.145041436 | 0.935787346 | -0.21263 | 0.998431 |

Supplementary Table 4: p values from Tukey HSD results comparing differences in raw and normalized energy-taxis diameter growth rates in pyruvate or succinate plates. Growth rates were obtained by dividing the raw or normalized diameter at 48h by the value at 24h. HSD test compared all pairs of strains, only comparisons to the wild-type are presented here. P values were computed using a 0.95 confidence value, those below 0.05 were taken as significant.

| Comparison | Pyruvate Normalized | Succinate Normalized | Pyruvate Raw | Succinate Raw |
|---|---|---|---|---|
| Wild-Type-Aer.g1 | 1 | 1 | 1 | 1 |
| Wild-Type-Aer.g1/Aer.g4 | 1 | 1 | 1 | 1 |
| Wild-Type-Aer.g2 | 1 | 1 | 0.999 | 1 |
| Wild-Type-Aer.g2/Aer.g1 | 0.997992 | 0.194316 | 0.96148 | 0.37878 |
| Wild-Type-Aer.g2/Aer.g1/Aer.g4 | 0.826779 | 1 | 0.332046 | 1 |
| Wild-Type-Aer.g2/Aer.g4 | 1 | 1 | 0.999447 | 1 |
| Wild-Type-Aer.g4 | 0.999305 | 0.999923 | 0.894703 | 0.999999 |
| Wild-Type-Aer2 | 0.791526 | 1 | 0.305462 | 1 |
| Wild-Type-Aer2/Aer.g1 | 0.990515 | 1 | 0.937616 | 1 |
| Wild-Type-Aer2/Aer.g1/Aer.g4 | 0.999856 | 1 | 0.99263 | 1 |
| Wild-Type-Aer2/Aer.g2 | 1 | 1 | 1 | 1 |
| Wild-Type-Aer2/Aer.g2/Aer.g1 | 0.776794 | 0.994759 | 0.290882 | 0.997895 |
| Wild-Type-Aer2/Aer.g2/Aer.g1/Aer.g4 | 1 | 1 | 1 | 1 |
| Wild-Type-Aer2/Aer.g2/Aer.g4 | 1 | 0.995619 | 1 | 0.999439 |
| Wild-Type-Aer2/Aer.g4 | 1 | 0.99969 | 1 | 0.999984 |
| Wild-Type-Aer2/CttP | 1 | 0.516674 | 1 | 0.066568 |
| Wild-Type-Aer2/CttP/Aer.g2/Aer.g1/Aer.g4 | 1 | 1 | 1 | 1 |
| Wild-Type-Aer2/CttP/Aer.g2/Aer.g4 | 1 | 0.999829 | 1 | 1 |
| Wild-Type-CheA | **0** | **0** | **0** | **0** |
| Wild-Type-CttP | 1 | 0.990882 | 1 | 0.996631 |
| Wild-Type-CttP/Aer.g1 | 0.866421 | 0.259124 | 0.070324 | **0.039548** |
| Wild-Type-CttP/Aer.g2 | **0.03075** | 1 | 0.416662 | 1 |
| Wild-Type-CttP/Aer.g2/Aer.g1/Aer.g4 | 1 | 0.999082 | 1 | 0.712737 |
| Wild-Type-CttP/Aer.g2/Aer.g4 | 1 | 1 | 0.999329 | 0.986805 |
| Wild-Type-CttP/Aer.g4 | 0.926885 | 1 | 0.999998 | 0.939527 |

Supplementary Table 5: p values from Tukey HSD results comparing differences in energy-taxis diameters, diameter changes and diameter growth rates in pyruvate or succinate plates for complementation strains. 5del indicates deletion of Aer2/CttP/Aer.g2/Aer.g1/Aer.g4. Growth rates were obtained by dividing the raw or normalized diameter at 48h by the value at 24h. Amount of growth was obtained by subtracting the 24h diameter from the value at 48h. HSD test compared all pairs of strains. P values were computed using a 0.95 confidence value, those below 0.05 were taken as significant (bold).

| Parameter | Time | Comparison | Succinate Difference | Succinate Adjusted p Value | Pyruvate Difference | Pyruvate Adjusted p Value |
|---|---|---|---|---|---|---|
| Diameter | 24 | 5del_342-5del_33 | -57.607 | 0.920 | -59.515 | 0.644 |
| | | wt_Aer.g1-5del_Aer.g1 | 254.831 | 0.157 | 119.757 | 0.204 |
| | | wt_342-5del_Aer.g1 | 149.687 | 0.463 | 93.301 | 0.346 |
| | | wt_Aer.g1-5del_342 | 312.438 | 0.089 | 179.272 | 0.067 |
| | | wt_342-5del_342 | 207.294 | 0.256 | 152.816 | 0.108 |
| | | wt_342-wt_Aer.g1 | -105.144 | 0.691 | -26.456 | 0.943 |
| Diameter | 48 | 5del_342-5del_Aer.g1 | -264.812 | 0.196 | -171.301 | 0.755 |
| | | wt_Aer.g1-5del_Aer.g1 | 624.510 | **0.014** | 425.043 | 0.199 |
| | | wt_342-5del_Aer.g1 | 207.487 | 0.331 | 309.349 | 0.382 |
| | | wt_Aer.g1-5del_342 | 889.321 | **0.004** | 596.343 | 0.079 |
| | | wt_342-5del_342 | 472.299 | **0.036** | 480.649 | 0.146 |
| | | wt_342-wt_Aer.g1 | -417.023 | 0.054 | -115.694 | 0.900 |
| Growth | NA | 5del_342-5del_Aer.g1 | -207.205 | **0.015** | -111.786 | 0.824 |
| | | wt_Aer.g1-5del_Aer.g1 | 369.679 | **0.002** | 305.286 | 0.228 |
| | | wt_342-5del_Aer.g1 | 57.800 | 0.461 | 216.048 | 0.441 |
| | | wt_Aer.g1-5del_342 | 576.884 | **0.000** | 417.072 | 0.102 |
| | | wt_342-5del_342 | 265.005 | **0.006** | 327.834 | 0.193 |
| | | wt_342-wt_Aer.g1 | -311.879 | **0.003** | -89.238 | 0.897 |
| Rate | NA | 5del_342-5del_Aer.g1 | -0.422 | 0.265 | -0.235 | 0.745 |
| | | wt_Aer.g1-5del_Aer.g1 | 0.134 | 0.892 | 0.430 | 0.362 |
| | | wt_342-5del_Aer.g1 | -0.224 | 0.674 | 0.284 | 0.639 |
| | | wt_Aer.g1-5del_342 | 0.556 | 0.136 | 0.665 | 0.136 |
| | | wt_342-5del_342 | 0.198 | 0.743 | 0.519 | 0.249 |
| | | wt_342-wt_Aer.g1 | -0.358 | 0.365 | -0.146 | 0.915 |

Supplementary Table 6: Plasmids used for energy-taxis experiments in *Pseudomonas pseudoalcaligenes* KF707.

| Plasmid | Description | Reference |
|---|---|---|
| pRK2013 | KmR ori colE1 RK2-Mob$^{+}$ RK2-Tra$^{+}$ | (2) |
| pG19II | Gm$^{R}$, *sacB*, *lacZ*, cloning vector, conjugative plasmid | (3) |
| pG19II-*Δaer-2* | Gm$^{R}$, *sacB*, *lacZ*, *aer-2* deletion construct | This Study |
| pG19II-*ΔcttP* | Gm$^{R}$, *sacB*, *lacZ*, *cttP* deletion construct | This Study |
| pG19II-*Δaer.g1* | Gm$^{R}$, *sacB*, *lacZ*, *aer.g1* deletion construct | This Study |
| pG19II-*Δaer.g2* | Gm$^{R}$, *sacB*, *lacZ*, *aer.g2* deletion construct | This Study |
| pG19II-*Δaer.g4* | Gm$^{R}$, *sacB*, *lacZ*, *aer.g4* deletion construct | This Study |
| pSEVA324 | Cm$^{R}$, pR01600/ColE1, *lacZα*-pUC19 | (4) |
| pSEVA324-*aer.g1* | Cm$^{R}$, pSEVA342 with *aer.g1* in the MCS | This Study |

Supplementary References


1. **Schultz J**, **Copley RR**, **Doerks T**, **Ponting CP**, **Bork P**. 2000. SMART: a web-based tool for the study of genetically mobile domains. Nucleic Acids Res **28**:231–4.

2. **Figurski DH**, **Helinski DR**. 1979. Replication of an origin-containing derivative of plasmid RK2 dependent on a plasmid function provided in trans. Proc Natl Acad Sci U S A **76**:1648–52.

3. **Maseda H**, **Sawada I**, **Saito K**, **Uchiyama H**, **Nakae T**, **Nomura N**. 2004. Enhancement of the mexAB-oprM efflux pump expression by a quorum-sensing autoinducer and its cancellation by a regulator, MexT, of the mexEF-oprN efflux pump operon in Pseudomonas aeruginosa. Antimicrob Agents Chemother **48**:1320–8.

4. **Silva-Rocha R**, **Martínez-García E**, **Calles B**, **Chavarría M**, **Arce-Rodríguez A**, **De Las Heras A**, **Páez-Espino AD**, **Durante-Rodríguez G**, **Kim J**, **Nikel PI**, **Platero R**, **De Lorenzo V**. 2013. The Standard European Vector Architecture (SEVA): A coherent platform for the analysis and deployment of complex prokaryotic phenotypes. Nucleic Acids Res **41**:D666–D675.
